# Three-dimensional cilia structures from animals’ closest unicellular relatives, the Choanoflagellates

**DOI:** 10.1101/2022.02.24.481817

**Authors:** Justine M. Pinskey, Adhya Lagisetty, Long Gui, Nhan Phan, Evan Reetz, Gang Fu, Daniela Nicastro

## Abstract

In most eukaryotic organisms, cilia perform a variety of life-sustaining roles related to environmental sensing and motility. Cryo-electron microscopy has provided considerable insight into the morphology and function of ciliary structures, but studies have been limited to less than a dozen of the millions of known eukaryotic species. Ultrastructural information is particularly lacking for unicellular organisms in the opisthokont clade, leaving a sizeable gap in our understanding of cilia evolution between unicellular species and multicellular metazoans (animals). Choanoflagellates are important aquatic heterotrophs, uniquely positioned within the opisthokonts as the metazoans’ closest living unicellular relatives. We performed cryo-focused ion beam milling and cryo-electron tomography on cilia from the choanoflagellate species *Salpingoeca rosetta.* We show that the axonemal dyneins, radial spokes, and central pair complex in *S. rosetta* more closely resemble metazoan structures than those of unicellular organisms from other suprakingdoms. In addition, we describe unique features of *S. rosetta* cilia, including microtubule holes, microtubule inner proteins, and the ciliary vane: a fine, net-like extension that has been notoriously difficult to visualize using other methods. Furthermore, we report barb-like structures of unknown function on the extracellular surface of the ciliary membrane. Together, our findings provide new insights into choanoflagellate biology and cilia evolution between unicellular and multicellular opisthokonts.

## Introduction

Most eukaryotic organisms possess motile cilia, which perform a variety of functions necessary for their survival. All major branches of the eukaryotic tree of life contain ciliated representatives, strongly suggesting the presence of one or more cilia in the last eukaryotic common ancestor (LECA) (Cavalier-Smith, 2002; Mitchell, 2004, 2007) (Figure 1). The vast majority of eukaryotic life consists of unicellular organisms with one or more cilia, which aid in motility, feeding, avoiding predators, and sensing the environment (Burki, 2014; Mitchell, 2007). Multicellular eukaryotes, including animals (metazoans), also rely on cilia for developmental signaling, mucosal clearance, feeding, and reproduction. The structure of motile cilia is quite complex and contains several hundred different proteins (Pazour et al., 2005). Yet the mutation of a single ciliary protein can result in severe ciliary assembly or motility defects, which can lead to death or disease, including human ciliopathies (Reiter and Leroux, 2017).

**Figure 1.**
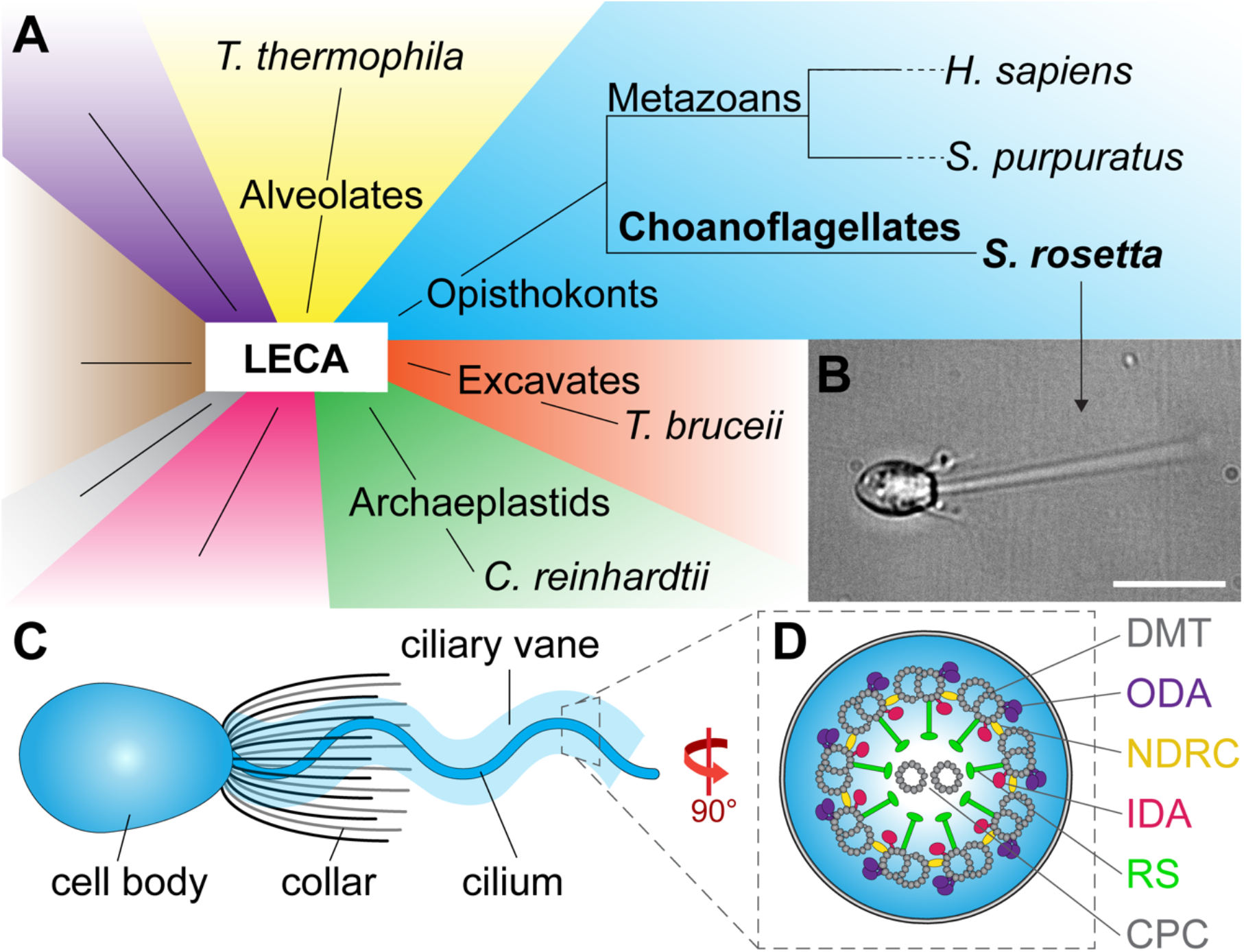
Phylogeny and ciliary features of the choanoflagellate *S. rosetta*. **A**) Phylogenetic tree showing major eukaryotic suprakingdoms (colored) stemming from the last common Eukaryotic ancestor (LECA). Suprakingdoms with representatives that have been imaged using cryo-ET are labeled (i.e. Alveolates, Opisthokonts, Excavates, and Archaeplastids) with example species. Choanoflagellates are part of the Opisthokont branch and form a sister group with metazoans, having shared a last common unicellular ancestor more than 600 million years ago. Whereas metazoans are multicellular animals, the choanoflagellates remained unicellular/colonial. **B**) Fixed *Salpingoeca rosetta* cell (a marine choanoflagellate). A short movie of an *S. rosetta* cell swimming and additional images of selected *S. rosetta* cell types can be found in Figure 1 supplements 1 and 2, respectively. **C**) Overview cartoon of the choanoflagellate cell architecture, including the cell body and the ring of actin-based collar tentacles surrounding a single motile cilium with a ciliary vane. **D**) Cross-sectional diagram of the choanoflagellate cilium indicating known ciliary components. Cross section in this figure and throughout the paper are viewed from proximal towards the distal tip of the flagellum, and longitudinal sections are shown with proximal on the left unless otherwise indicated. Labels: CPC, central pair complex; DMT, doublet microtubule; IDA and ODA, inner and outer dynein arm; N-DRC, nexin-dynein regulatory complex; RS, radial spoke. Scale bar: 10 µm (B).

Though the overall architecture of motile cilia is conserved, ciliary protein structures, accessory features, and regulatory complexes show some divergence throughout evolution. Most motile cilia contain a ring of nine outer doublet microtubules (DMTs) with a pair of central singlet microtubules, often referred to as the “9+2” arrangement (Fawcett, 1954) (Figure 1D), although exceptions exist, such as the vertebrate nodal cilia (9+0), eel sperm flagella (9+0) and rabbit posterior notochord cilia (9+4) (Takeda and Narita, 2012). The axonemal core in motile cilia contains 96-nm repeat units with two rows of dyneins, the outer and inner dynein arms (ODAs, IDAs), regulatory complexes like the nexin-dynein regulatory complex (N-DRC) and radial spokes (RSs) (Figure 1D), and the central pair complex (CPC), again with some exceptions (Porter and Sale, 2000; Smith and Yang, 2004). Despite these broad commonalities, ultrastructural studies have shown differences in the morphology of ciliary protein complexes (Lin et al., 2014). Motile cilia also exhibit a variety of beating patterns including helical, planar, base to tip, tip to base, or reversible (Blake and Sleigh, 1974), and can be outfitted with an assortment of accessory structures, including mastigoneme hairs, paraflagellar rods, fibrous sheaths, outer dense fibers and accessory microtubules (de Souza and Souto-Padron, 1980; Hyams, 1982; Irons and Clermont, 1982a, b; Mencarelli et al., 2008; Nakamura et al., 1996; Portman and Gull, 2010; Yubuki et al., 2016).

Our understanding of ciliary ultrastructure and evolution is continually expanding through application of new technologies. Historically, much of our knowledge of ciliary architecture from diverse species has been based on conventional light and electron microscopy studies, which are inherently limited by detection limits and preservation artifacts. Protein sequence comparisons have also yielded important insights, particularly into dynein evolution in eukaryotic cilia (Kollmar, 2016), although this required manual annotation of thousands of genes from hundreds of species, not particularly sustainable for examining hundreds of ciliary proteins. Furthermore, sequence comparisons are limited in their ability to predict protein morphology, localization, and interactions. As a result, our knowledge of detailed ciliary morphology, function, and evolution has remained restricted. Advances in cryo-electron tomography (cryo-ET) have enabled visualization of native ciliary structures with unparalleled resolution, enhancing our ability to compare ciliary morphology across species and make inferences about their evolution and function, although cilia from less than a dozen species have currently been examined using cryo-ET (Carbajal-Gonzalez et al., 2013; Fu et al., 2018; Lin et al., 2012a; Lin et al., 2012b; Lin and Nicastro, 2018; Lin et al., 2014; Nicastro et al., 2011; Pigino et al., 2012).

Motile cilia from several multicellular animals (metazoans) have been studied using cryo-ET, but high-resolution structural information is lacking for unicellular organisms in the same opisthokont clade, preventing structural comparison between metazoans and their close unicellular relatives. Cryo-ET studies of unicellular species from other suprakingdoms, such as archaeplastida (e.g. *Chlamydomonas*), alveolata (e.g. *Tetrahymena*) and excavata (e.g. *Trypanosoma*) (Figure 1A), have revealed significant morphological differences between unicellular and multicellular motile cilia, including dynein number and arrangement, CPC shape and microtubule inner proteins, and radial spoke head morphology (Carbajal-Gonzalez et al., 2013; Imhof et al., 2019; Lin et al., 2014; Pigino et al., 2011; Pigino et al., 2012). However, these clades are phylogenetically quite distant from metazoans, raising questions about when and how these differences arose throughout the evolutionary timescale (Figure 1A). Choanoflagellates are unicellular (or colonial) organisms within the opisthokont branch that share a last common ancestor with metazoans (the urchoanozoan) more than 600 million years ago (Carr et al., 2008; King, 2004; Ruiz-Trillo et al., 2008; Steenkamp et al., 2006). Because of their unique phylogenetic position, choanoflagellates provide important information on the origin and evolution of multicellular organisms (King, 2004). Though low-resolution features of choanoflagellate cilia have been described (Hibberd, 1975; Karpov, 2016; Leadbeater, 2015), detailed molecular structures remain unexamined.

Here, we use cryo-focused ion beam milling (cryo-FIB) and cryo-ET to investigate the cilium and other structures in the ciliary region of the marine choanoflagellate species *S. rosetta*. Our tomographic reconstructions and 3D averages suggest that choanoflagellates and their metazoan relatives share similar morphology and arrangement of ciliary dyneins and their regulators, suggesting that these features were already present in the two groups’ last common ancestor. Similarly, the *S. rosetta* CPC strongly resembles that of *S. purpuratus* sperm flagella, with the exception of a partially reduced C1d projection. In contrast, however, we also observed ciliary features that appear to be unique to Choanoflagellates, such as previously unseen gaps and microtubule inner proteins (MIPs) in the DMTs, the ciliary vane, which is a fine mesh of intertwined filaments extending bilaterally from the ciliary membrane, and barb-like structures, which protrude from the extracellular surface of the ciliary membrane. These findings expand our understanding of choanoflagellate biology and provide insights into the evolution of motile ciliary structures within the opisthokont branch.

## Results

*S. rosetta* contain a single cilium which extends from the cell body and is surrounded by a ring of 25-36 actin-based collar tentacles (Figure 1, Figure 1-figure supplements 1 and 2) (Dayel et al., 2011). As microbial filter feeders, choanoflagellates use the planar beat of their cilium to generate both cell motility and microcurrents, which enable them to more easily engulf bacterial prey (Pettitt et al., 2002). The overall structure of the choanoflagellate cilium has been previously studied using light and conventional electron microscopy techniques, revealing a 9+2 axonemal microtubule arrangement and a basal body that is surrounded by a microtubule rootlet structure (Karpov, 2016; Karpov and Leadbeater, 1998). We sought to visualize molecular structures within and surrounding the *S. rosetta* cilium with improved resolution enabled by technical advances in cryo-FIB milling and cryo-ET imaging (Marko et al., 2007; McIntosh et al., 2005).

### Cryo-ET and subtomogram averaging facilitate high-resolution analyses of the S. rosetta cilium

*S. rosetta* can transition between several cell types, including single-celled slow and fast swimmers, doublets, chains, rosettes, and thecate cells that attach to substrates through a secreted basal process (examples in Figure 1-figure supplement 2) (Dayel et al., 2011). We rapidly froze starved choanoflagellate singlet cells in their slow- and fast-swimming morphological states. During plunge-freezing, areas close to the cell body were embedded in relatively thick ice (>500 nm); therefore, we used cryo-FIB milling to generate ∼150-200 nm thin lamellae (sections) of the plunge-frozen cells before cryo-ET imaging (Figure 2, A-F). In one cryo-FIB lamella, we captured part of the cell body with actin-filled collar filaments extending outward and the proximal region of the cilium (Figure 2, D). From this lamella, we were able to record sequential cryo-tomograms along the ciliary length (Figure 2, E-F). Within the reconstruction of the basal apparatus, we observe part of the basal body and the surrounding MTOC ring of dense material from which the lateral rootlet microtubules radiate outwards (Figure 2E) (Karpov, 2016; Pozdnyakov et al., 2017). We observe multiple collar bases and many vesicles distributed throughout the basal pole (Figure 2E). In addition, the ciliary vane filaments were clearly visible on two opposite sides of the cilium and extended to the edges of the imaging area (∼3 µm) (Figure 2D, F). Farther away from the cell body, the ice was sufficiently thin to perform cryo-ET imaging directly on the plunge-frozen cilia (Figure 2, G and H). The motile cilium, actin-based collar tentacles and thin vane filament were readily visible in the 3D reconstructions (Figure 2, E-H).

**Figure 2.**
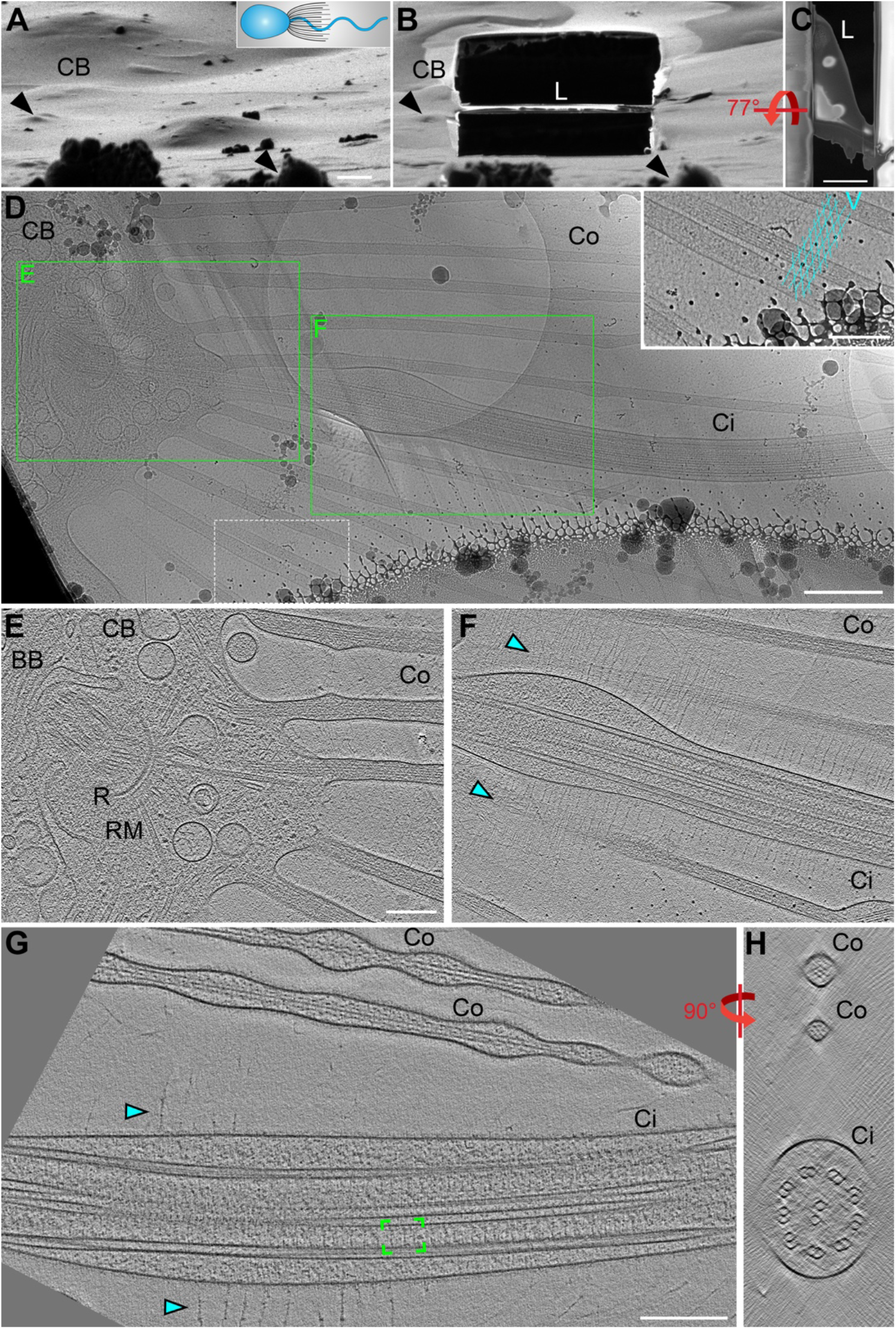
Cryo-FIB milling and cryo-ET enable visualization of ciliary structures. **A-C**) Choanoflagellate cell before (**A**) and after (**B**, **C**) cryo-FIB milling, as viewed by the ion beam (**A**, **B**) and the electron beam (**C**). The cartoon denotes the cell’s orientation, with cell body (CB) to the left. In (**A**, **B**), the stage with the EM grid is tilted to approximately 13 degrees relative to the ion beam for milling. A perpendicular top view of the cryo-FIB lamella is shown in (**C**). Black arrowheads denote surface features in the ice to serve as landmarks for positional orientation. Note: the lamella (L) that includes the cilium appears very low relative to the cell body, but that is a visual illusion caused by the tilt and the several micron thick sputter/GIS-layer on top of the ice layer. **D**) Overview map of the milled cilium (Ci), with green boxes indicating the positions of two sequential tomograms that were recorded from this lamella, shown in (**E** and **F**). The area within the white dashed line is magnified as an inset in the upper right corner, highlighting the regular meshwork of vane filaments which extend past the edges of the map. **E**-**F**) Tomographic slices emphasizing the basal body (BB) and collar filaments (Co) (**E**) and the proximal region of the cilium (**F**). Cyan arrowheads denote vane filaments in (**F**, **G**). **G**-**H**) Example of a whole cell (i.e. not cryo-FIB milled) tomographic reconstruction of a *S. rosetta* cilium in longitudinal (**G**) and cross-sectional (**H**) views. Green brackets indicate a single 96-nm axonemal repeat, thousands of which were averaged together to generate the subtomogram averages shown in Figure 3. Other labels: R, ring of dense material (MTOC); RM, rootlet microtubules. Scale bars: 2 μm (C); 1 μm (A, applies also to B); 500 nm (D); 200 nm (D inset; E, applies also to F; G, applies also to H).

To better resolve the molecular details of the *S. rosetta* cilium, we performed subtomogram averaging of >7500 axonemal repeats (96 nm length) that were extracted from 54 cryo-tomograms (Figure 2G, green brackets; Figure 3), which yielded an average with 2.2 nm resolution (0.5 FSC criterion) (Figure 3-figure supplement 1; Table 1). With this resolution, we observe that the axonemal repeats of *S. rosetta* cilia contain outer dynein arms with two motor domains each, the double-headed I1 (f) inner dynein complex, and six single-headed inner dynein arms, *a*, *b*, *c*, *e*, *g*, and *d* (Figure 3). Most motile cilia contain three radial spokes per axonemal repeat (RS1-RS3), which project from the A-tubule toward the CPC, and regulate ciliary motility through poorly understood signaling mechanisms (Zhu et al., 2017). The *S. rosetta* cilium also contains three full-length radial spokes per axonemal repeat (Figure 3, C, C’, E, and F) with somewhat variable radial spoke head positions, causing them to blur-out slightly in the averages (Figure 3, A and C). This positional flexibility was likely because intact *S. rosetta* cells were frozen while their cilia were actively beating. To improve the resolution of the radial spoke heads, we performed local alignments focused on each of the three head domains (Figure 3C’). Similar to human cilia, the shape of the *S. rosetta* radial spoke heads resemble narrow ice skates (Figure 3, C’ and F; Figure 7), rather than the broad radial spoke head morphology of other unicellular species like *Chlamydomonas* and *Tetrahymena* (Figure 7) (Barber et al., 2012; Lin et al., 2014; Pigino et al., 2011; Pigino et al., 2012; Poghosyan et al., 2020). The head domains of *S. rosetta* RS1 and RS2 are separated from one another, whereas those of RS2 and RS3 are connected (Figure 3, C, E, and F).

**Figure 3.**
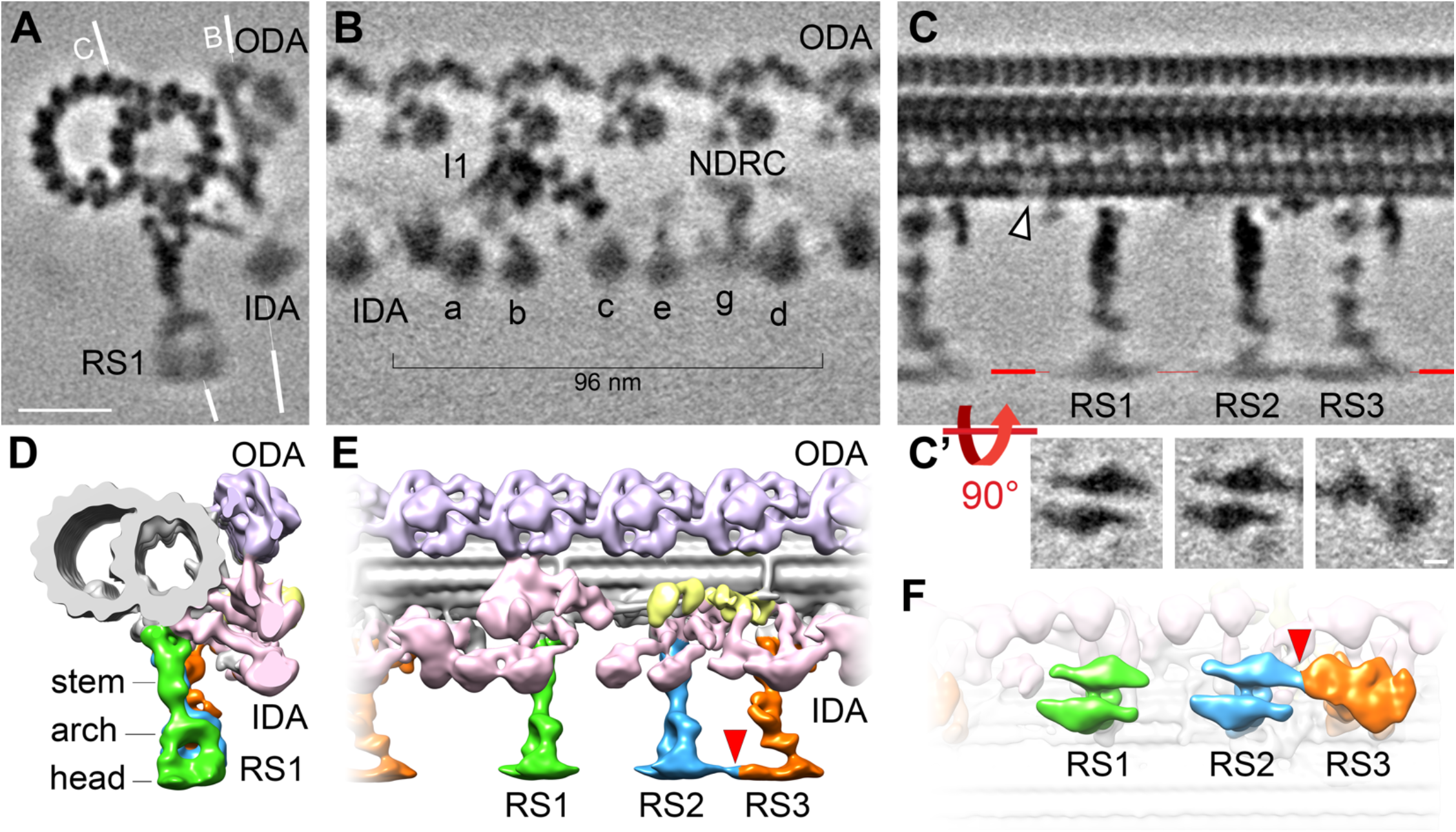
Cryo-ET of native choanoflagellate cilia reveals structural features of the 96-nm axonemal repeat. **A-C’**) Cross- sectional (**A**) and longitudinal (**B**, **C**) sections through the subtomogram average of the doublet microtubule; the white lines in (**A**) indicate the positions of the longitudinal sections shown in (**B**, **C**). Major doublet associated complexes are labelled, such as the outer and inner dynein arms (ODA, lavender; IDA, a-e, g, pink), the I1 dynein (I1, pink), the nexin-dynein regulatory complex (NDRC, yellow), and the radial spokes (RS1, 2, and 3, green, blue, and orange, respectively). The radial spoke heads were blurred in the averages due to positional heterogeneity, therefore we performed local alignment refinements for each radial spoke head, which are displayed viewed from the bottom in (**C’**). The white arrowhead in (**C**) denotes a hole in the A-tubule, which appears blurred because the hole was not present in all averaged repeats (see classification in Figure 4). Resolution information and tomogram/particle numbers for the subtomogram average displayed in panels (**A**-**C**) are in Figure 3-figure supplement 1 and Table 1. Figure 3-figure supplement 2 includes additional information on DMT-specific features. **D-F**) Isosurface rendering of the averaged *S. rosetta* 96-nm axonemal repeat shown in cross-sectional (**D**), longitudinal (**E**), and bottom (**F**) views. Scale bars: 20 nm (A, applies to A-C); 5 nm (applies to all panels in C’).

**Figure 7.**
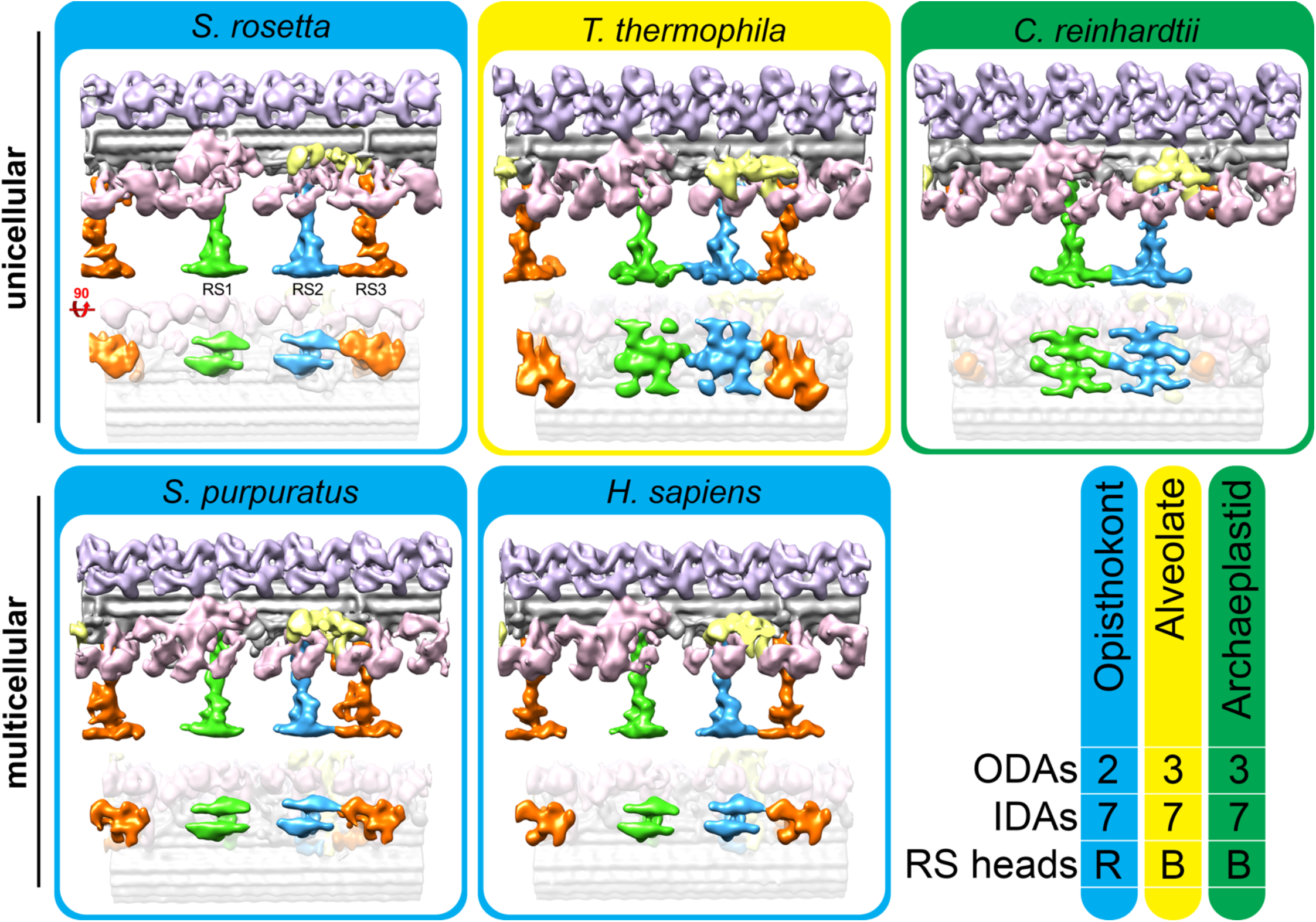
The ciliary structures of the unicellular choanoflagellate more closely resembles those of multicellular opisthokonts than unicellular organisms from other suprakingdoms. Isosurface renderings of the 96-nm ciliary repeats from unicellular species (top row) vs. multicellular animals (metazoans; bottom row). The summary on the bottom right highlights that the cilia of opisthokonts, including the unicellular/colonial *S. rosetta*, contain two outer dynein arms (ODAs) and reduced (R) radial spoke (RS) heads, whereas Alveolates and Archaeplastids contain three ODAs and broad-shaped (B) RS heads. Each organism contained one double-headed and seven single-headed inner dynein arms (IDAs). The averaged axonemal structures from species other than *S. rosetta* were previously published in (Lin et al., 2014).

**Table 1.** Summary of data included in this study

| Specimen | Tomograms included | Averaged repeats | Resolution at 0.5 Fourier shell correlation criterion (nm) | Resolution at 0.143 Fourier shell correlation criterion (nm) | Used in Figure(s) |
| --- | --- | --- | --- | --- | --- |
| *S. rosetta* slow/fast swimmers<sup>a</sup> | 54 | 7584 | 2.2 | 1.8 | 3, 4, 7 |
| Central Pair Complex<sup>b</sup> | 28 | 1323 | 2.5 | 2.2 | 5 |
| Barb structures (with 4-fold symmetry)<sup>c</sup> | 17 | 600 | 2.5 | 1.9 | 6 |

<sup>a</sup> Resolution was estimated at the base of RS1 from a 64 voxel subvolume;
<sup>b</sup> Resolution was estimated at the central portion of the barb from a 64 voxel subvolume;
<sup>c</sup> Resolution was estimated at C1a from a 32 voxel subvolume.

Doublet-specific averages allowed us to identify the conserved bridge structure between DMTs 5 and 6 (Afzelius, 1959; Lin et al., 2012b), thus orienting the doublet numbers DMT1-9 within each reconstructed cilium (Figure 3-figure supplement 2). Similar to *S. purpuratus* (Lin et al., 2012b), DMT 5 in *S. rosetta* cilia lacks ODAs and a subset of IDAs (b, c, and e), replaced instead by the o-SUB5-6 and i-SUB5-6 structures (Figure 3-figure supplement 2; DMT5, green and orange arrowheads). We also observed a unique connection between the A-tubule and the base of IDA c on DMT 9, with a smaller partial density near the base of the A-tubule on DMT 1 (Figure 3-figure supplement 2, mustard arrowheads). An additional DMT-specific density corresponds to a dangling structure near the A2 hole described in Figure 4, which is present or partially present on DMTs 1, 2, 5, 6, and 7, but only weakly present or absent on DMTs 3, 4, 8, and 9 (Figure 3-figure supplement 2, navy blue arrowheads).

**Figure 4.**
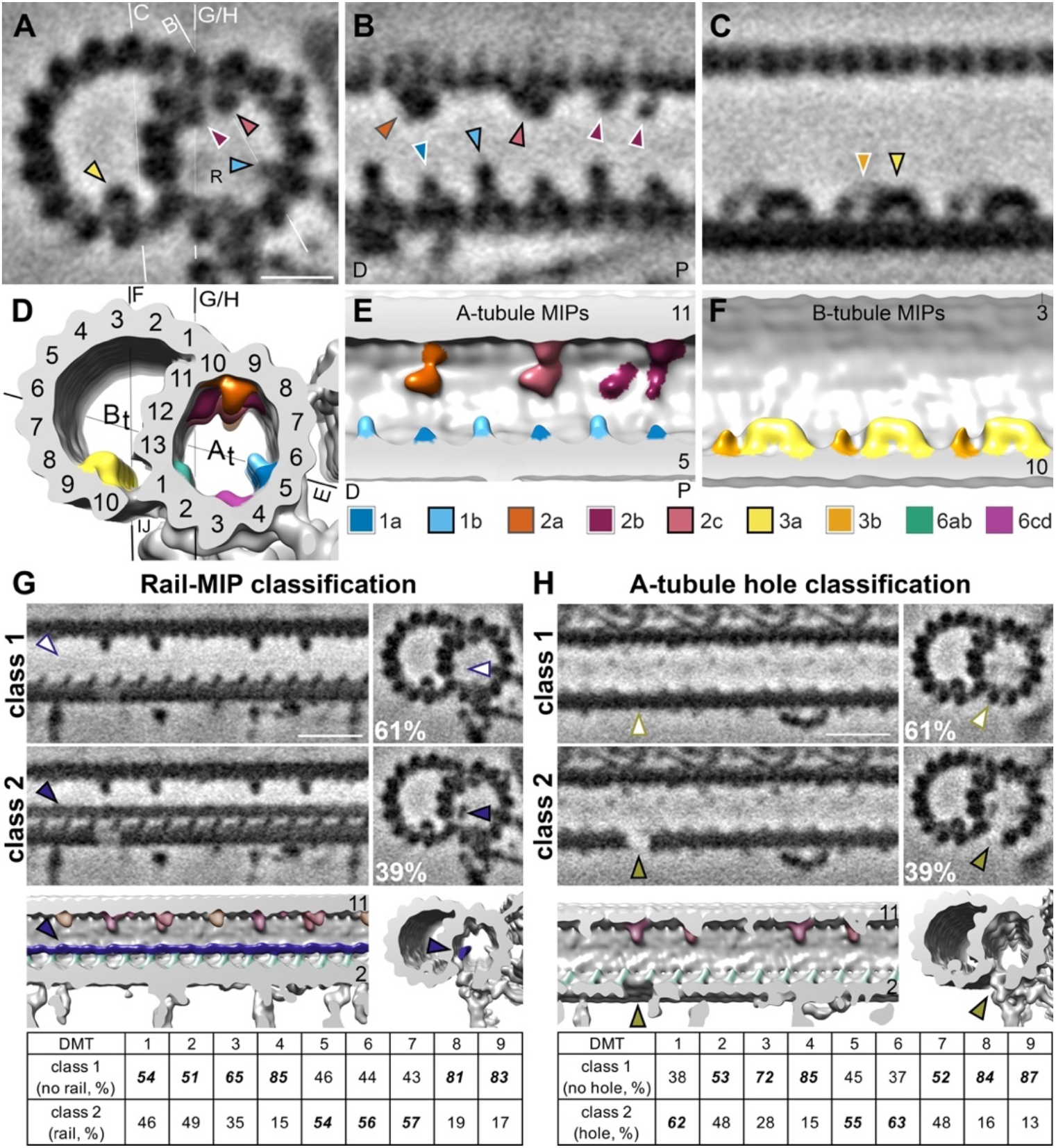
*S. rosetta* microtubule doublets contain unique holes and MIPs. **A**-**F**). Tomographic slices (**A**-**C**) and isosurface renderings (**D**-**F**) of the subtomogram average of *S. rosetta* doublet microtubules shown in cross-section at the level of RS1 (**A**, **D**), longitudinal section of the A-tubule (**B**, **E**), and longitudinal section of the B-tubule (**C**, **F**); the white lines in (**A**) indicate the positions of the longitudinal views shown in (**B**, **C**, **G**, **H**), and the black lines in (**D**) indicate the viewing positions of the isosurface renderings shown in (**E**, **F**, **G**, **H**). Note: panels (**B**, **E**) portray the distal side of the cilium to the left (distal and proximal indicated by D and P, respectively). MIPs (and arrowheads indicating MIPs) colored as indicated in the legend below panels (**E/F**). Because the MIPs repeat with a periodicity of 48 nm or less, only a 48 nm long segment of the 96-nm axonemal unit is shown. **G**, **H**) Classification analyses focused on the region with the newly identified rail-MIP (**G**) and A-tubule hole (**H**) show structural heterogeneity, specifically absence and presence of these features in subsets of the axonemal repeats. Classes 1 (top rows) lack the rail-MIP or A-tubule hole (empty arrowheads), whereas classes 2 (bottom rows) contain the rail-MIP (navy blue arrowhead) or A-tubule hole (olive arrowhead), respectively; percentages of repeats out of 7584 averaged particles are indicated for each class. The isosurface renderings highlight the position of the rail-MIP (navy blue) between protofilaments A1 and A13, adjacent to MIP 6ab (jade) (**G**), and of the A-tubule hole in protofilament A2 (**H**). The quantification below shows the doublet-specific distribution of the classes – for example, 54% of repeats from DMT 1 separated into class 1 (no rail-MIP), while 46% were in class 2 (rail-MIP). Note: the A-tubule hole also appears mildly stronger in the rail-MIP class 2 (**G**), but a weakening of the density is also present in the rail-MIP class 1 due to only a partial overlap of the rail-MIP and A-tubule hole distribution. Overall, both features show some doublet-specific distribution, with the rail-MIP preferentially being present on doublets 5-7 and the A-tubule hole on doublets 1, 5, and 6. Figure 4-figure supplement 1 provides a more detailed representation of the rail-MIP and A-tubule hole classification results for each studied tomogram; the analyses show that the rail-MIP is either mostly present or mostly absent depending on the tomogram, suggesting a possible difference between (tomograms from) the proximal or distal ciliary region, whereas the A-tubule hole classes are more evenly distributed. Figure 4-figure supplement 2 shows a FIB-milled cilium near the cell body, indicating the presence of the rail-MIP in the proximal cilium. Figure 4- figure supplement 3 shows two additional holes in the *S. rosetta* inner junction. Scale bars: 10 nm (A, applies also to B, C); 20 nm (G, applies to all other images in the panel); 20 nm (H, applies to all other images in the panel).

### S. rosetta doublet microtubules show conserved and unique features

Microtubule inner proteins (MIPs) are regularly distributed proteins that attach to the luminal side of the microtubule wall in cilia (Ichikawa et al., 2017; Kirima and Oiwa, 2018; Maheshwari et al., 2015; Nicastro et al., 2011; Nicastro et al., 2006), and in other hyperstable microtubule species, including subpellicular microtubules in apicomplexan parasites (Wang et al., 2021) and the ventral disc microtubules of *Giardia* (Schwartz et al., 2012). Many of the ciliary MIPs are highly conserved between species (Ichikawa et al., 2017; Khalifa et al., 2020; Ma et al., 2019; Maheshwari et al., 2015; Nicastro et al., 2011; Nicastro et al., 2006; Song et al., 2020), but some species-specific MIP features have also been reported, such as the *Chlamydomonas* beak-MIP (Dymek et al., 2019; Hoops and Witman, 1983), the *T. brucei*- specific B2, B4, B5, ponticulus MIPs, snake-MIP, ring MIP, and Ring Associated MIP (RAM) (Imhof et al., 2019), and a connection of MIP3 to the mid-partition in *Tetrahymena* (Li et al., 2022). Based on their locations and periodicities along the 96-nm repeat, we identified many conserved MIPs within *S. rosetta* ciliary doublet microtubules, including MIPs 1a, 1b, 2a, 2b, 2c, 3a, 3b, and 6a-d (Figure 4, A-F). However, we also observed some differences, e.g. MIP1a is typically longer than MIP1b in other species (Song et al., 2020), but in *S. rosetta* MIP1a is shorter than MIP1b (Figure 4, A-B, D-E). Furthermore, we identified a previously unobserved ∼3.5 nm wide filamentous MIP, named rail-MIP here, which runs along the length of the A- tubule near protofilament A13 and seems to connect to MIP 6ab (Figure 4A and G-class 2). The electron density of this rail-MIP was reduced in the average of all axonemal repeats (Figures 3A, 4A), suggesting its presence on only a subset of repeat units. To further explore this heterogeneity, we performed automated classification analyses (Heumann et al., 2011) focused on the rail-MIP by applying a mask around the region of interest and using principle component analyses to sort the results into identifiable features. Indeed, these analyses revealed that the rail- MIP was not ubiquitously present: only 39% of all averaged axonemal repeat units contained the rail-MIP, and its presence was enriched in DMTs 5-7 (Figure 4G, class 2 and table), as compared to doublets 1-4 and 8-9, which mostly lacked the rail-MIP (Figure 4G, class 1 and table). The rail-MIP distribution varied between tomograms: about half of the tomograms contained prevalent rails concentrated in the microtubules stated above, whereas the other half of ciliary reconstructions contained fewer, scattered rail-MIPs (Figure 4-figure supplement 1). This asymmetric distribution does not appear to correlate with any other observed features (such as presence of vane, barbs, collar filaments, or IFT particles), but the rail-MIP is present in a cryo- FIB-milled lamella containing the proximal region of the cilium (Figure 4-figure supplement 2), suggesting that its distribution could be related to the location of the tomogram along the length of the cilium (proximal vs. distal).

Ciliary DMTs of most (wild-type) species described so far by cryo-ET display one ∼4 nm long hole per axonemal repeat in the inner junction between protofilaments A1 and B10; the only described exception is the *T. brucei* ciliary DMTs that has an additional (more proximal) inner junction hole per repeat (Imhof et al., 2019). We and others have shown that one PACRG- subunit is missing near the N-DRC base-plate from the FAP20-PACRG inner junction filament (Dymek et al., 2019; Ma et al., 2019; Nicastro et al., 2011). In addition to this inner junction-hole near the N-DRC, *S. rosetta* cilia contain two additional inner junction-holes, one near the base of RS1, and one near the base of RS3 (Figure 4-figure supplement 3). Distances between the previously-reported N-DRC-related inner junction-hole and the additional proximal and distal hole are ∼32 and ∼16 nm, respectively, suggesting that they could represent additional PACRG subunit losses, given the 8-nm periodicity of the FAP20-PACRG repeat (Dymek et al., 2019).

Notably, the location of the proximal inner junction hole in *S. rosetta* does not correspond to the proximal inner junction-hole in *T. brucei*, which is ∼48 nm proximal of the previously-reported N-DRC-related inner junction-hole (Imhof et al., 2019). *S. rosetta* ciliary DMTs also exhibit a (so far unique) ∼6.5 nm long gap in protofilament A2 of the A-tubule between RS3 and RS1 from the next axonemal repeat unit, likely due to a missing tubulin dimer (Figure 4H, class 2; Figure 4-figure supplement 3). Like the heterogeneous rail-MIP, the electron density in the position of the A2-hole was reduced but not completely missing in the average of all axonemal repeats (Figure 3C, Figure 4-figure supplement 3), suggesting its presence on only a subset of the repeats. Our classification analyses revealed that the A2-hole is present in ∼39% of repeats, including over 50% of repeats from DMTs 1, 5, and 6 and with lower frequencies in the other DMTs (Figure 4H table: class 2). Unlike the rail-MIP, the distribution of the A2-hole across tomograms did not cluster, instead appearing relatively evenly scattered throughout different tomograms (Figure 4-figure supplement 1). Although the DMT-specificity between the rail-MIP and A2-hole somewhat overlapped (presence in DMTs 5 and 6), there was only a mild correlation between these two unique DMT features within repeat units, as evidenced by the hole appearing mildly stronger in class 2 containing the rail-MIP, but still somewhat present in class 1 without the rail-MIP (Figures 4G-H, Figure 4-figure supplement 1).

### The S. rosetta central pair complex shows overall conserved features with some reductions

The CPC forms the central core of the axoneme in most motile cilia and consists of two singlet microtubules (C1 and C2) that are surrounded by a specific set of projections (Carbajal-Gonzalez et al., 2013). In some organisms, the CPC is fixed in its orientation relative to the doublet microtubules, whereas in others, such as *Chlamydomonas*, it twists within the axoneme (Omoto et al., 1999). Like other opisthokonts, the *S. rosetta* CPC has a relatively fixed orientation, and the plane that contains both CPC microtubules is roughly parallel to the 5-6 bridge (Figure 5-figure supplement 1). This orientation is consistent with *S. rosetta’s* cilia having a planar, sinusoidal waveform (Dayel et al., 2011; Dayel and King, 2014) with an amplitude (beating direction) that is perpendicular to the CPC and 5-6-bridge planes.

**Figure 5.**
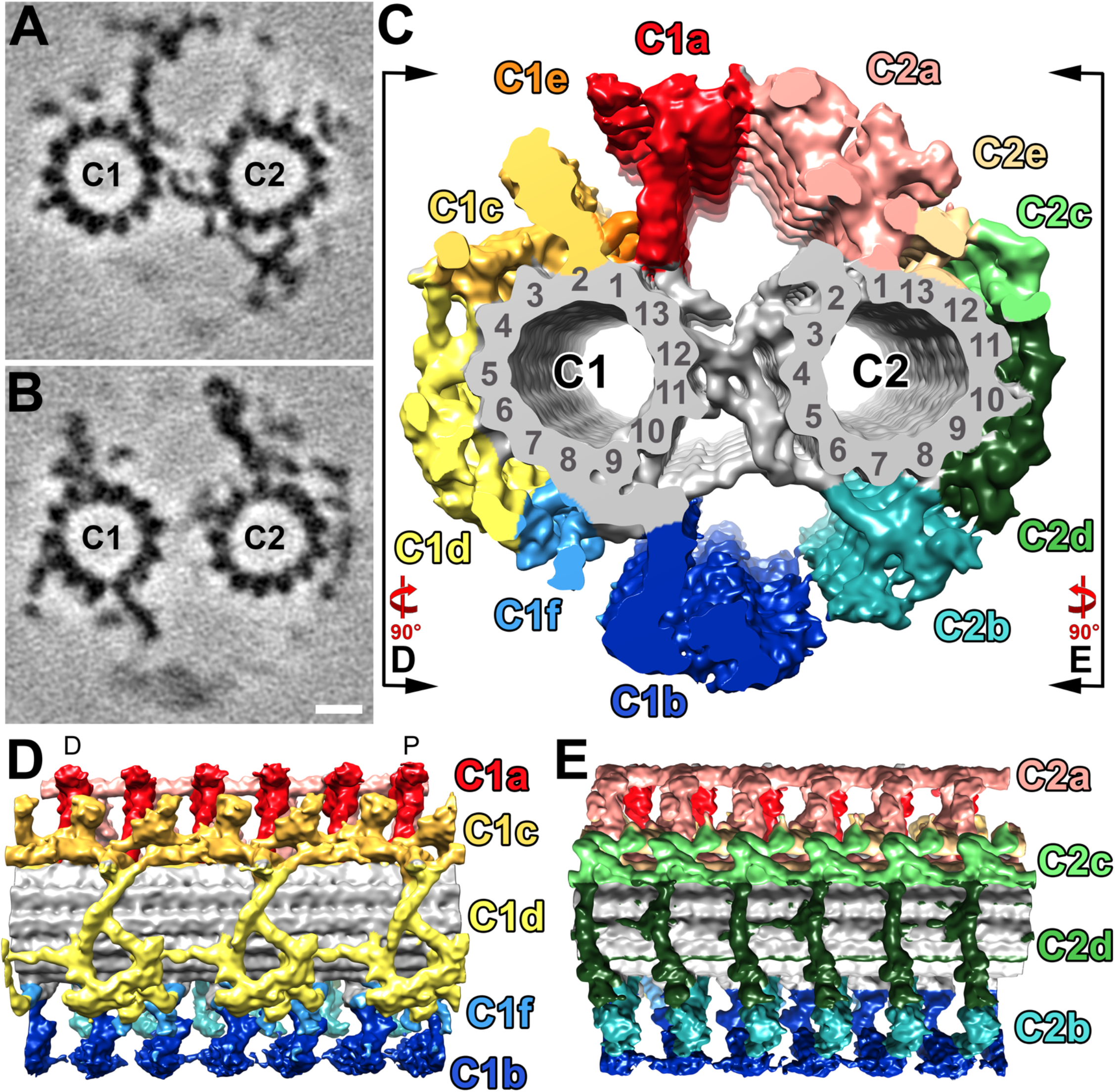
Structural features of the *S. rosetta* central pair complex. **A**-**C**) Tomographic slices at two different positions (**A**, **B**) and isosurface rendering (**C**) of the averaged *S. rosetta* CPC. The tomographic slices highlight the following CPC projections: C1a, C2b, and the central bridge (**A**), and C1b-c, C2a, and C2c-e (**B**), respectively. Coloring of the projections in (**C**) follows (Carbajal-Conzalez et. al., 2013). Averages were generated using 1323 particles from 28 different tomograms (Resolution information in Figure 3-figure supplement 1 and Table 1). Black lines and rotation arrows in (**C**) indicate the viewing directions shown in (**D**, **E**). **D**-**E**) Isosurface renderings show longitudinal side-views of the averaged *S. rosetta* CPC. Note: panel (**D**) is oriented with the distal side of the cilium to the left (distal and proximal indicated by D and P, respectively). The orientation of the CPC in relation to the 5-6 bridge, vane, and barb structures is shown in Figure 5-figure supplement 1. Additional species comparisons are provided in Figure 5-figure supplement 2. Scale bar: 10 nm (B, applies also to A).

To better resolve the molecular details of the *S. rosetta* CPC, we performed subtomogram averaging of >1300 repeats (32 nm length) that were extracted from 28 cryo-tomograms (selected based on best image signal-to-noise ratio) (Figure 5), which yielded an average with 2.5 nm resolution (0.5 FSC criterion) (Figure 3-figure supplement 1, Table 1). The *S. rosetta* CPC contains all major projections described in other organisms with the previously described longitudinal periodicities of 16 nm (C1a, b; C2a, b, c, d, and e) and 32 nm (C1c, d, e, f) (Figure 5, Figure 5-figure supplement 2) (Carbajal-Gonzalez et al., 2013; Fu et al., 2019). In many ways, the *S. rosetta* CPC strongly resembles that of sea urchin sperm flagella (Carbajal-Gonzalez et al., 2013; Fu et al., 2019): both lack the C2 MIP, have a small C1e projection, and exhibit prominent connections between the C1a and C2a projections, as compared to the *Chlamydomonas* CPC (Figure 5, Figure 5-figure supplement 2). One unique feature of the *S. rosetta* CPC, however, is the partial reduction of the C1d protein network, specifically it seems that FAP54 is lacking (Han et al., 2022), which exposes a larger area of the C1 microtubule wall (Figure 5, Figure 5-figure supplement 2).

### The ciliary vane is a bilayer of mesh-like, extracellular filaments

The ciliary vane is a “mysterious” structure on either side of the proximal area of the choanoflagellate cilium that has long escaped electron microscopists using traditional methods (Leadbeater, 2006). Computer modeling predicts that a vane is necessary to generate fluid motion that would allow bacteria to be phagocytosed by the choanoflagellate collar filaments and cell body (Nielsen et al., 2017), but the presence of a vane structure itself has only been shown for a few freshwater species, including *Codosiga botrytis*, *Salipingoeca frequentissima*, *Monosiga brevicolis*, and *Salpingoeca amphoridium*, and at low resolution (Hibberd, 1975; Leadbeater, 2006; Mah et al., 2014). In contrast to previous proposals that the vane might be limited to freshwater species, we clearly observed vane filaments extending laterally from either side of the ciliary membrane in the marine *S. rosetta*, both in the cryo-FIB lamella of the proximal ciliary region (Figure 2, D and F; Figure 4-figure supplement 2), as well as more distally, where the cilium is embedded in ice thin enough to be directly imaged using cryo-ET (Figure 2G, Figure 6). The vane originates at the base of the cilium, and its edges often extend the entire width of (and maybe beyond) our tomograms and lamella, which are ∼1.2 µm (tomograms) to ∼3 µm (lamella) wide (Figure 2D-G, Figure 4-figure supplement 2, Figure 6, Figure 6-figure supplement 1). The plane containing the vane varies somewhat in relation to the CPC/sub-5-6 planes, but in most tomograms, it appears to be oriented roughly parallel to the CPC within an angle of up to ∼30° between the planes (Figure 5-figure supplement 1). This orientation is consistent with the vane’s predicted function in the pumping mechanism of these filter feeders (Nielsen et al., 2017), as the vane would be positioned to experience maximum hydrodynamic drag to push liquid and prey close to the collar filaments. We observed only two exceptions (out of 54 tomograms) where the vane plane was almost perpendicular to the CPC/sub5-6 plane (Figure 5-figure supplement 1I).

**Figure 6.**
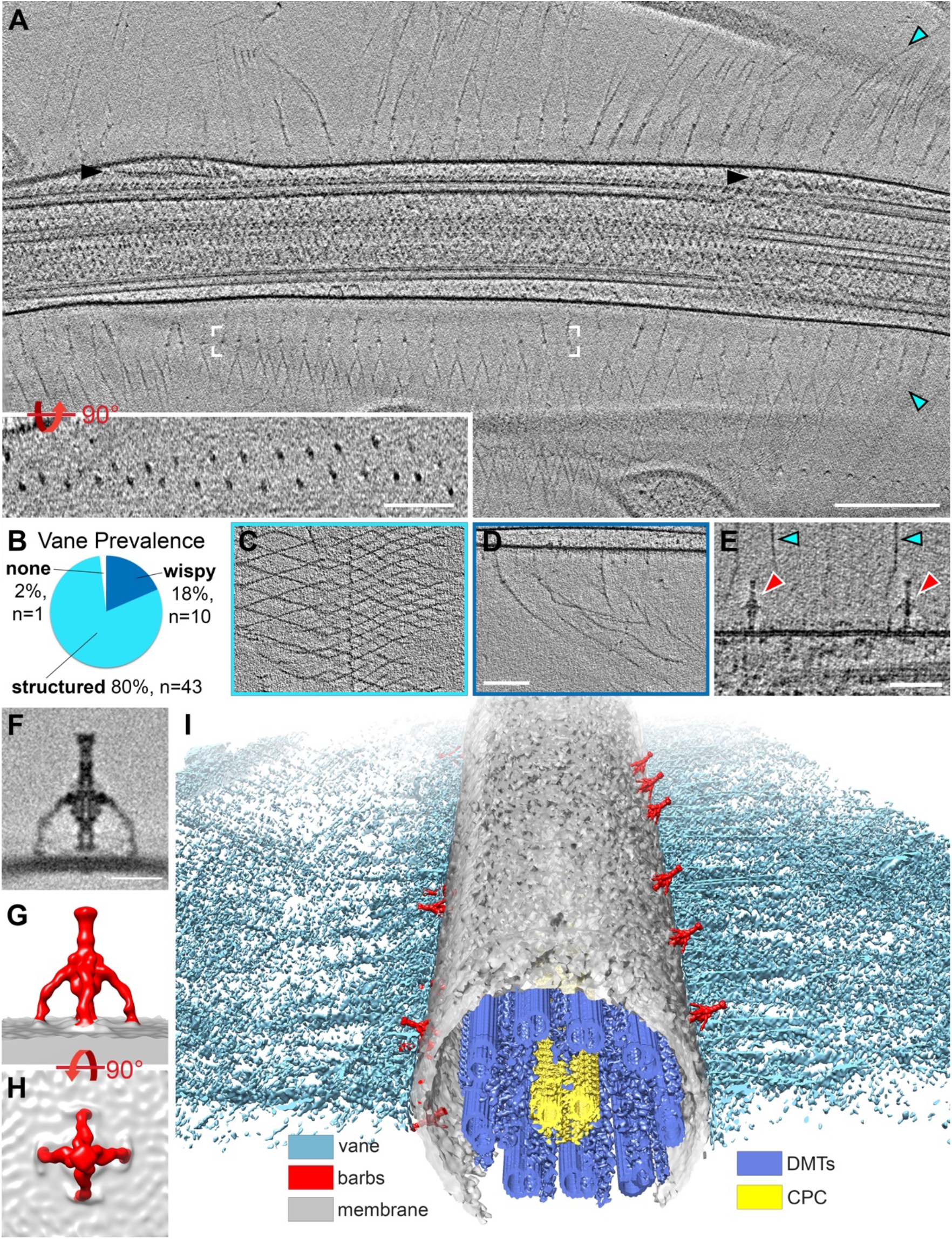
*S. rosetta* cells have a ciliary vane and adjacent barb structures. **A**) Tomographic slice through a representative cilium showing bilateral vane filaments (cyan arrowheads) extending from the ciliary membrane. Black arrowheads denote IFT trains. White brackets mark the region shown in the rotated inset, which shows that the vane is a bilayer of thin filaments with semi-regular spacing. **B**-**D**) Most tomograms contained vane filaments with regular patterning (“structured”, light blue color in (**B**), example tomographic slice shown in (**C**)), whereas a smaller proportion contained only individual “wispy” hairs (dark blue color in (**B**), example tomographic slice shown in (**D**)). Of the 54 tomograms included in our analyses, only one did not contain vane filaments. **E**) Tomographic slice showing two barb structures (red arrowheads), approximately 50 nm in height that protrude from the ciliary membrane near the plane of the vane filaments (cyan arrowheads). **F-H**) Tomographic slice (**F**) and isosurface renderings (**G**, **H**) show side (**F**, **G**) and top (**H**) views of the averaged barb structures (red, 4x symmetrized, 600 particles). **I**) Compiled isosurface rendering of the *S. rosetta* cilium, indicating positions of the vane (cyan) and barbs (red) relative to the ciliary membrane (gray); doublet microtubules (DMTs) and central pair complex (CPC) are shown in blue and yellow, respectively. Figure 6-figure supplement 1 shows vane filaments and barbs on the plasma membrane within the ciliary pocket, but not on the surface of *E. pacifica* (bacterial prey). Scale bars: 200 nm (A and inset); 100 nm (D, applies also to C); 50 nm (E); 20 nm (F).

Our data suggest that the *S. rosetta* ciliary vane is composed of two sheets of thin filaments on either side of the cilium, which extend from the ciliary membrane for approximately 80 nm before they split and attach to neighboring filaments to form diamond-shaped meshes (Figure 6, A-D). One or two rows of “nodes” are visible near the ciliary membrane where the filaments first split apart (Figure 2F, Figure 6A, Figure 4-figure supplement 2). Because our reconstructions are three dimensional, we can view these nodes rotated 90 degrees around the x-axis, which clearly shows that the two sheets of vane filaments and the nodes are separated by ∼45 nm (Figure 6A, inset). The nodes of the vane filaments are offset from one another so that the vertices of the diamonds from one sheet are centered within the diamonds from the overlaying sheet when viewed from top down (Figure 6, A and C). Consistent with previous reports, we observed some vane filaments that were highly organized and interconnected, whereas others appeared to be wispy, individual filaments (Figure 6, B-D). Vane filaments were apparent in all but one of the 54 tomograms we analyzed, with most tomograms exhibiting at least partial mesh-like structures (Figure 6 B-D). We also observed vane filaments basal to the ciliary membrane within the ciliary pocket (Figure 6-figure supplement 1). Vane filaments were not observed on the surface of the bacterial prey, *E. pacifica*, which were co-cultured with *S. rosetta* as a food source (Figure 6-figure supplement 1).

### Previously undescribed barb structures protrude from the S. rosetta ciliary membrane

In regions of the ciliary membrane adjacent to where the vane filaments protrude, we also observed previously undescribed barb-like membrane complexes, hereafter denoted as “barbs”, that extend ∼50 nm from the extracellular surface of the ciliary membrane (Figure 6, E-I). To better resolve the molecular details of these barbs, we performed subtomogram averaging and applied four-fold symmetry resulting in 600 averaged particles that yielded an average with 2.5 nm resolution (0.5 FSC criterion) (Figure 6 F-I; Table 1). The barbs consist of a top knob and a central rod with a wider mid-body, from which four arms protrude to connect to the membrane (Figure 6, E-I). Although the top knob of the barb resembles the size of the nodes of the ciliary vane (∼7 nm diameter), barbs were not observed at the base of each vane filament. Instead, they appeared loosely in two rows, one above and one below the vane bilayer on either side of the cilium, irregularly spaced along the cilium length (Figure 6I). The number of barbs varied greatly between tomograms (which show ∼2 µm ciliary length), ranging from 0-22 barbs, but their location was consistently near the base of the vane filaments (Figure 6I, Figure 5-figure supplement 1). Though a lumen is visible throughout parts of the central rod, the cavity/channel does not appear to be continuous, and the base of the rod seems only weakly connected to the membrane, if at all. We also observed several barb-like structures within the ciliary pocket (Figure 6-figure supplement 1). The overall shape of the barbs resembles head-less bacteriophages or bacterial secretion needles, but barb structures were not observed on the surface of 3D reconstructed *E. pacifica* bacterial cells, which were co-cultured with *S. rosetta* as a food source (Figure 6-figure supplement 1).

## Discussion

Cilia are hallmarks of eukaryotic cells, dating back to the LECA (Cavalier-Smith, 2002; Mitchell, 2004). Ciliary defects disrupt many important cellular functions and cause a variety of diseases in humans, collectively known as ciliopathies (Reiter and Leroux, 2017). Though detailed structural information is continually emerging, little is known about high-resolution cilia ultrastructure from diverse species. Furthermore, how cilia ultrastructure may have changed from unicellular to multicellular animals remains unexplored. Within the opisthokont clade, high-resolution structures of motile cilia have been published for several multicellular metazoans, including sea urchins, zebrafish, mouse, pig, horse, and humans (Leung et al., 2021; Lin et al., 2014; Nicastro et al., 2006; Yamaguchi et al., 2018; Zhao et al., 2021; Zheng et al., 2021).

However, no unicellular opisthokonts have been studied at similar resolution. Choanoflagellates are the closest living unicellular relatives to metazoans, and choanoflagellate studies have led to countless insights about the origins of multicellularity and the evolution of multicellular structures and processes (King, 2004). Here, we present high-resolution cilia structures from the choanoflagellate species *S. rosetta*, providing insights into the structural basis for choanoflagellate motility and a foundation to explore the evolution of cilia ultrastructure between uni- and multi-cellular opisthokonts.

### Ciliary evolution and the last common ancestor between choanoflagellates and metazoans

Though eukaryotic cilia are highly conserved overall, ultrastructural studies have revealed interesting dichotomies between unicellular and multicellular specimens. Most unicellular species contain three outer dynein heads (i.e. three dynein heavy chains) per ODA, whereas cilia from multicellular species typically contain only two (Lin et al., 2014; Pigino et al., 2012) (Figure 7). This is consistent with comparative genomic data suggesting that the third outer dynein heavy chain was lost in metazoans and – most likely independently – in excavates (Kollmar, 2016). In addition, unicellular organisms like *Tetrahymena* and *Chlamydomonas* have broad RS head structures and connections between all three radial spoke heads, whereas metazoan RS head structures are narrow, especially RS1 and RS2 are reduced to a pair of thin blades, and RS1 and RS2 are clearly separated from one another (Figure 7) (Lin et al., 2014).

Furthermore, the CPC orientation is fixed in metazoans, with stable connections between C1a and C2a and a reduced C1e projection (Carbajal-Gonzalez et al., 2013). Here, we find that the structure and organization of the dyneins, radial spokes, and the CPC observed for metazoan cilia are consistent with those of the unicellular *S. rosetta*. This suggests that these changes were likely present in the urchoanozoan (the phylogenetic group containing all Choanoflagellates and metazoan), pre-dating the transition to multicellularity.

Why might these ultrastructural changes have occurred? We can speculate that bulkier ciliary structures like the third outer dynein head, broad radial spoke heads, and larger CPC projections may have been lost to accommodate additional molecular components in the cilia that were related to the increased sensing and signaling functions between cells in multicellular organisms. Although free-swimming unicellular eukaryotes also signal through their motile cilia (sensing light, chemical environmental cues, and mechanosensory stimuli) cilia have adapted many additional signaling functions and molecules in animals, including T2R, TRP/PKD channels, progesterone receptors, estrogen receptor-ß, interleukin-6 receptor, and Hedgehog (HH) pathway components (Bloodgood, 2010; Mitchell, 2007). Choanoflagellate cilia fall somewhere in between, lacking GPCRs, Arrestins, GRK’s, and HH molecules, but containing TRP channel-related proteins like ECV3A/B and Enkurin, suggesting the association of signaling molecules with the ciliary compartment prior to the animal origin (Sigg et al., 2017). Many choanoflagellate species spend part or all of their life cycle attached to substrates using carbohydrate-based theca structures or attached to one another in sheets or colonies (Dayel et al., 2011; Leadbeater, 2015). Could the reduced RS head structures, CPC projections, and dynein motors in *S. rosetta* be related to a shift away from prey capture and predator avoidance requiring a strong ciliary beat, towards increased signaling functions in a more stable and protected environment? Future studies might consider examining ciliary structures from earlier-branching opisthokonts as well as other sessile filter feeders to expand on these comparisons.

On the other hand, choanoflagellates can exist in free-swimming, single-cell states, and they must find food, avoid predators, and survive harsh aquatic environments like other unicellular organisms. How might they compensate for the decreased ciliary stability that may have resulted from the reduction of ciliary structures (i.e. RS heads, dynein motors, CPC)? We report a unique rail-MIP in *S. rosetta* cilia that runs the length of the A-tubule lumen. MIPs are thought to reinforce structural integrity of doublet microtubules (Owa et al., 2019), therefore a combination of the rail-MIP and A2-hole identified here may provide the strength and flexibility necessary to compensate for the loss of bulkier ciliary structures. The rail-MIP is found preferentially in specific doublets with a distribution that resembles the also asymmetric distribution of the beak-MIPs in the B-tubule of DMTs 1,5,6 of *Chlamydomonas* cilia (Nicastro et al., 2011). Thus, it is also possible that both the rail-MIP and beak-MIP provide mechanical support to compensate for the extra drag generated by the motion of the ciliary vane (choanoflagellates) or mastigonemes (*Chlamydomonas*) through the aqueous environment, given both structures’ wing-like positioning perpendicular to the beating direction of the flagella.

Likely, a combination of these factors has enabled choanoflagellates to reduce their dynein and radial spoke structures without losing their cell-propulsion function.

### Enhanced ciliary vane preservation reveals its detailed and unique morphology

The choanoflagellate ciliary vane has remained understudied due to technical challenges with fixation and visualization of this filigree structure. Plunge-freezing and cryo-ET overcomes these challenges, allowing us to study the vane in unprecedented detail, both confirming and extending previous observations and interspecies comparisons. The choanoflagellate ciliary vane has only been observed in a few freshwater species of the >125 known choanoflagellate species (Hibberd, 1975; Leadbeater, 2006; Mah et al., 2014). These reports describe the vane as a bilateral fringe composed of delicate perpendicular fibers of glycocalyx, occasionally with diagonal or longitudinal fibers, which extend approximately two-thirds of the flagellar length (Hibberd, 1975; Leadbeater, 2006; Mah et al., 2014). Our data are consistent with these reports and further resolve two layers of vane filaments on each side of the cilium, which split and connect with neighboring filaments and are offset to form a mesh-like appearance (Figure 2D,F,G, 6A-D, I). The wispy hairs we observe appear similar to the “partially disintegrated” vanes observed in *Monosiga* sp. (Hibberd, 1975), suggesting that they are perhaps not disintegrated but rather a common variation on vane structure.

As has been previously discussed (Hibberd, 1975), the structure of the choanoflagellate vane differs significantly from other flagellar appendages, including the hair-like mastigonemes from green algae like *Chlamydomonas* (Liu et al., 2020) or the tripartite hairs from golden algae like *Ochromonas* (Bouck, 1971). Both the size and arrangement of filaments is distinct, with algal mastigonemes comprised of two intertwined filaments with an overall diameter of ∼10 nm and organized as single or tripartite hairs (Liu et al., 2020), vs choanoflagellate filament diameters of ∼3.5 nm arranged as wispy hairs or meshed networks. In addition, we do not detect connections between the membrane anchor of the choanoflagellate vane filaments and the axonemal microtubules, in contrast to observations for *Chlamydomonas* and *Euglena* (Liu et al., 2020). Consistent with these morphological differences, we also did not detect homologues of the mastigoneme protein MST1 and membrane anchor PKD2 in the *S. rosetta* genome via BLAST search, suggesting that the vane differs from these flagellar appendages. Instead, it was proposed that the morphology of the choanoflagellate vane is similar to the bilateral, wing-like vane of sponge choanocyte flagella, which is anchored in the ciliary membrane without connections to the axonemal microtubules (Mehl and Reiswig, 1991). However, sponge choanocyte vanes appear to be narrower, denser, and more massive than choanoflagellate vanes, and they often connect laterally to the collar filaments, which we and others do not observe in *S. rosetta* (Brunet and King, 2017; Hibberd, 1975; Leadbeater, 2006; Mah et al., 2014; Mehl and Reiswig, 1991). Future studies on the ultrastructure of the sponge choanocyte ciliary vane, as well as the composition of both choanocyte and choanoflagellate vane filaments will further elucidate the extent of their similarity and provide insight as to their evolutionary relationship.

How do these delicate and intricate vane structures form, and what are they made of? Though our data do not directly address these questions, we do observe fibers that originate within the ciliary pocket, suggesting the possibility that the vane could be secreted from the cell membrane before being transported into the ciliary compartment (Figure 6-figure supplement 1). Though not present in our cryo-ET data of fast and slow swimmers, large, fiber-filled vesicles have been observed near the apical part of the choanoflagellate cells following division, presumably when the cilium and vane would be regenerating (Leadbeater, 2015). Intriguingly, glycosyltransferases such as those encoded by *jumble* and *couscous* localize to both the basal pole and the ciliary/collar base, and the collar base also stains positively for jacalin, *Lycopersicon esculentum* (tomato) lectin (LEL), and *Solanum tuberosum* (potato) lectin (STL), indicating the presence of carbohydrate chains made of Galß3GalNAc and GlcNAc2-4 near the choanoflagellate cilium (Wetzel et al., 2018). We performed an external digest with proteinase K, which failed to remove the ciliary vane (data not shown), supporting the hypothesis that the *S. rosetta* ciliary vane is carbohydrate or glycoprotein-based rather than proteinaceous, though additional study is necessary to further characterize the specific vane component(s).

### Barb structures and their possible functions

Adding to the ‘mysterious’ composition and function of the ciliary vane (Leadbeater, 2006), we have also identified previously undescribed barb structures attached to the *S. rosetta* ciliary membrane. At about 50 nm in height and with four “arms” and a central rod connecting to the ciliary membrane, the barbs’ function and composition present yet additional mysteries. Like the vane filaments, we find barb structures in both the ciliary membrane and the plasma membrane of the ciliary pocket (Figure 6-figure supplement 1). Despite of their resemblance with head-less bacteriophages or bacterial secretion needles, the barbs’ semi-regular distribution in two loose rows around the vane filaments, makes it unlikely that the barb structures originate from an external source – for instance the co-cultured *E. pacifica* bacterial prey. Consistently, we do not observe barb structures on the surface or inside of *E. pacifica* cells or floating in the medium (Figure 6-figure supplement 1). However, the *S. rosetta* genome has undergone extensive horizontal gene transfer from bacteria and other organisms (Matriano et al., 2021), so it is possible that the barbs could have an ancient bacterial origin. Indeed, the barb structures most closely resemble the size, shape, and general distribution of an uncharacterized bacterial complex in *Prosthecobacter debontii*, though the bacterial structures exhibit a 5-fold rather than 4-fold symmetry (Dobro et al., 2017).

The barbs’ location alongside the ciliary vane suggests that they could play a role in vane generation or maintenance. Indeed, the top knobs at the barbs’ distal ends resemble the size and shape of the nodes within the proximal vane, though the distance to the ciliary membrane in the vane base is roughly twice that of the barb height. Although beyond the scope of this study, finding cellular contexts in which barbs are increased or decreased, e.g. during states in which the cilium and vane are regenerating following cell division or artificially induced, could provide additional insight into the barb function.

With continually improving cryo-ET workflows and new techniques for genetic modification in choanoflagellates (Booth and King, 2020; Booth et al., 2018), there has never been a more exciting time for detailed ultrastructural and functional analyses in our closest unicellular relatives. This work raises many important questions, particularly regarding the role of ciliary structural changes and extracellular matrix components in choanoflagellate biology. Combining recent advances in opisthokont phylogeny with morphological and ultrastructural traits, we can better predict the nature of the last common ancestor of choanoflagellates and animals. Furthermore, because choanoflagellate cilia more closely resemble human cilia they may represent an attractive alternative to other protist model systems like *Chlamydomonas* and *Tetrahymena* for the study of human ciliopathies.

## Materials and Methods

### Choanoflagellate culture and cryo-preparation

*S. rosetta* co-cultured with *E. pacifica* was obtained from ATCC (PRA-390) and was cultured as previously described (King lab choanoflagellate handbook: https://kinglab.berkeley.edu/resources/). Briefly, cells were maintained in 5% Seawater Complete Media (SWC) diluted with artificial seawater (ASW), and cells swimming in the top half of the flask were passed 1:3 every 2-4 days.

Before freezing, cultures were scaled up to 100-400 mL, pelleted at 2000 x g (4°C, 10-15 min), resuspended in ASW and starved for 20-24 hours to reduce excess *E. pacifica*. Starved *S. rosetta* cells were then similarly pelleted and resuspended in a small volume of ASW (100 uL – 1 mL), and the concentrated cell sample was mixed 3:1 with 10 nm BSA-coated colloidal gold (Iancu et al., 2006) shortly before plunge-freezing. 4 µL of the mixture were pipetted onto a copper EM grid with holey carbon film (R2/2, 200 mesh, Quantifoil Micro Tools GmbH, Q43486) that had been freshly glow-discharged for 30 s at -30 mAmp. Samples were back-blotted for 1-3 s with Whatman filter paper (grade 1) to remove excess buffer and immediately plunge frozen into liquid ethane using a homemade plunge freezing device. Vitrified samples were mounted into either Autogrids with notched ring for FIB-milling or regular Autogrids for direct cryo-ET (Thermo Fisher Scientific) and stored in liquid nitrogen until used.

### Cryo-FIB milling

Autogrids with vitrified *S. rosetta* were cryogenically transferred into an Aquilos dual-beam FIB/SEM instrument (Thermo Fisher Scientific) equipped with a cryo-stage that was precooled to -185 °C. To protect the sample and enhance conductivity, layers of platinum were added to the grid surface (sputter-coater: 1 keV and 30 mA for 20s, gas injection system (GIS): pre-heated to 27 °C and deposited onto the sample for 5 s) (Schaffer et al., 2017). An overview image of the grid was generated in SEM mode, and cells suitable for cryo-FIB milling were identified using the Maps software. For milling, the cryo-stage was tilted to a shallow angle of 11-18 degrees between the EM grid and the gallium ion beam. Cryo-FIB milling was performed using a 30 keV gallium ion beam with currents of 30 pA for initial bulk milling and thinning, and 10 pA for final polishing, resulting in ∼150 nm thick, self-supporting lamella. SEM imaging at 3 keV and 25 pA was used to monitor the milling process.

### Cryo-ET imaging

Vitrified samples and cryo-FIB milled lamella were imaged using a 300 keV Titan Krios transmission electron microscope (Thermo Fisher Scientific) equipped with a Bioquantum post-column energy filter (Gatan) used in zero-loss mode with a 20 eV slit width and a Volta Phase Plate with -0.5 μm defocus (Danev et al., 2014). The SerialEM microscope control software (Mastronarde, 2005) was used to operate the Krios and record dose-symmetric tilt series (Hagen et al., 2017) from -60° to +60° tilt with 2° increments. Tilt series images were collected using a K3 Summit direct electron detector (Gatan) at 26,000x magnification and under low-dose conditions and in counting mode (for each tilt series image: 10 frames, 0.05 s per frame, dose rate of ∼28 e/pixel/s, frames were recorded in super-res mode and then binned by 2, resulting in a pixel size of 3.15 Å). The cumulative electron dose per tilt series was limited to 100 e^-^/Å^2^.

### Data processing

Preprocessing and 3D reconstruction were performed using the IMOD software package (Kremer et al., 1996). K3 frames were dose-weighted and motion-corrected using Motioncorr2. The tilt series images for whole cell reconstructions were aligned using the 10 nm gold nanoparticles as fiducials. Images for lamella reconstructions were aligned either fiducial-less using patch-tracking in IMOD or using dark features (e.g. from the sputter coat or embedded Gallium) as fiducials. 3D reconstructions were calculated using weighted back-projection.

Tomograms were excluded from further analysis if they contained compressed cilia or were damaged by non-vitreous ice. Subtomogram averaging was performed as previously described using the PEET program (Nicastro et al., 2006). Initial averages if the barb structures suggested symmetry, thus four-fold symmetry was applied during the final steps of subtomogram averaging. To sort particles (i.e. axonemal repeats) with and without rail-MIP and/or A2 hole, soft-edged masks were applied around those features and unsupervised classification analyses built into the PEET program (Heumann et al., 2011) was performed to calculate class averages (Figure 4, G and H). For clearer views of the radial spoke heads, local alignment refinement was performed focused on each individual RS heads. Isosurface renderings were generated using the UCSF Chimera package software (Pettersen et al., 2004). Resolution of the 96-nm repeat, CPC, and barb particle were estimated at the base of RS1, C1a projection, and particle center, respectively, using the Fourier shell correlation method with a criterion of 0.5 (Figure 3-figure supplement 1, Table 1). Tomographic slices (without subtomogram averaging) were denoised for better presentation using either non-linear anisotropic diffusion (Figure 2E-H and Figure 6A, C-E) or a weighted median filter (smooth filter in IMOD) (Figure 4-figure supplement 2, Figure 5- figure supplement 2, Figure 6-figure supplement 1).

### Light microscopy

For live-cell imaging, 5-10 µL *S. rosetta* cultures were pipetted directly onto superfrost plus glass slides (Fisherbrand) between two thin streaks of petroleum jelly (applied using a 22- gauge needle with syringe), over which an 18-mm circle glass coverslip (Fisherbrand) was gently suspended to create vertical space for the cells to move freely. Brightfield fast time-lapse series were acquired on a Nikon Eclipse Lvdia-N microscope using a 40x 0.75 NA Plan Fluor objective and the Nikon Elements software, later converted to .mov using Fiji. For culture images, *S. rosetta* cells were fixed with 2% glutaraldehyde (Sigma-Aldrich, Germany) for 10 min, and 5-10 µL were transferred to glass slides, covered with a glass coverslip, and sealed with clear nail polish. DIC images were collected on an inverted Nikon Eclipse Ti microscope using a 60x 1.4- NA Plan Apochromat oil objective, an Orca-Fusion digital camera (Hamamatsu), and the Nikon Elements software.

## Acknowledgements

The authors would like to acknowledge Drs. Julie Pfeiffer and Arielle Woznica (UT Southwestern Medical Center) for their generous assistance sharing reagents and knowledge of choanoflagellate cell culture, and Drs. Jeffrey Woodruff and Maralice Connaci-Sorell for sharing equipment, reagents, and advice. We are also grateful to Dr. Daniel Stoddard, Jose Martinez, and Eric Zhang of the Cryo-EM Facility at UT Southwestern Medical Center, as well as current and previous Nicastro lab members, for their ongoing assistance and support. Cryo-EM data were collected at the UT Southwestern Medical Center Cryo-Electron Microscopy Facility, which is supported in part by the CPRIT Core Facility Support Award RP170644. This study was funded by the following grants: National Institutes of Health grants R01GM083122 (to D.N.) and F32 GM137470 (to J.M.P.), and a Cancer Prevention and Research Institute of Texas (CPRIT) grant RR140082 (to D.N.). This research was also supported in part by the computational resources provided by the BioHPC supercomputing facility located in the Lyda Hill Department of Bioinformatics, UT Southwestern Medical Center. Cryo-ET data for the 96-nm ciliary repeat average, central pair complex average, and barb average have been deposited to the EM Data Bank under accession codes EMD-26204, EMD-26209, and EMD-26210, respectively.

## Author Contributions

J.M.P contributed to conceptualization, methodology, validation, formal analysis, investigation, data curation, writing original draft, writing reviewing and editing, visualization, supervision, project administration, and funding acquisition. A.L. contributed to validation, formal analysis, investigation, and data curation. L.G. contributed to formal analysis, data curation, and visualization. N.P. and E.R. contributed to methodology. G.F. contributed to conceptualization and investigation. D.N. contributed to conceptualization, methodology, validation, writing reviewing and editing, visualization, supervision, project administration, and funding acquisition.

---

**Figure 1-figure supplement 1.**
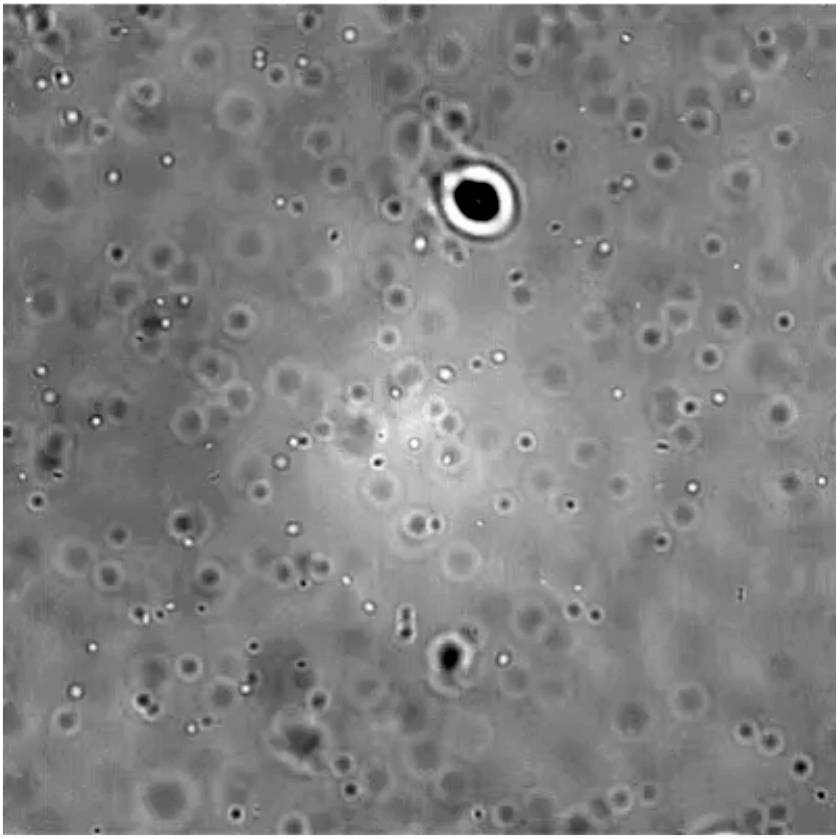
(video, see file attached). Choanoflagellate cell swimming in culture imaged by light microscopy at 40x magnification. Propelled by its single cilium, a slow swimming *S. rosetta* cell (dark/larger oval) navigates through a field of bacterial prey (small circles). Captured bacteria also line the ring of collar filaments at the base of the cell body. A recently divided doublet cell enters the field of view from the lower right near the end of the video.

**Figure 1-figure supplement 2.**
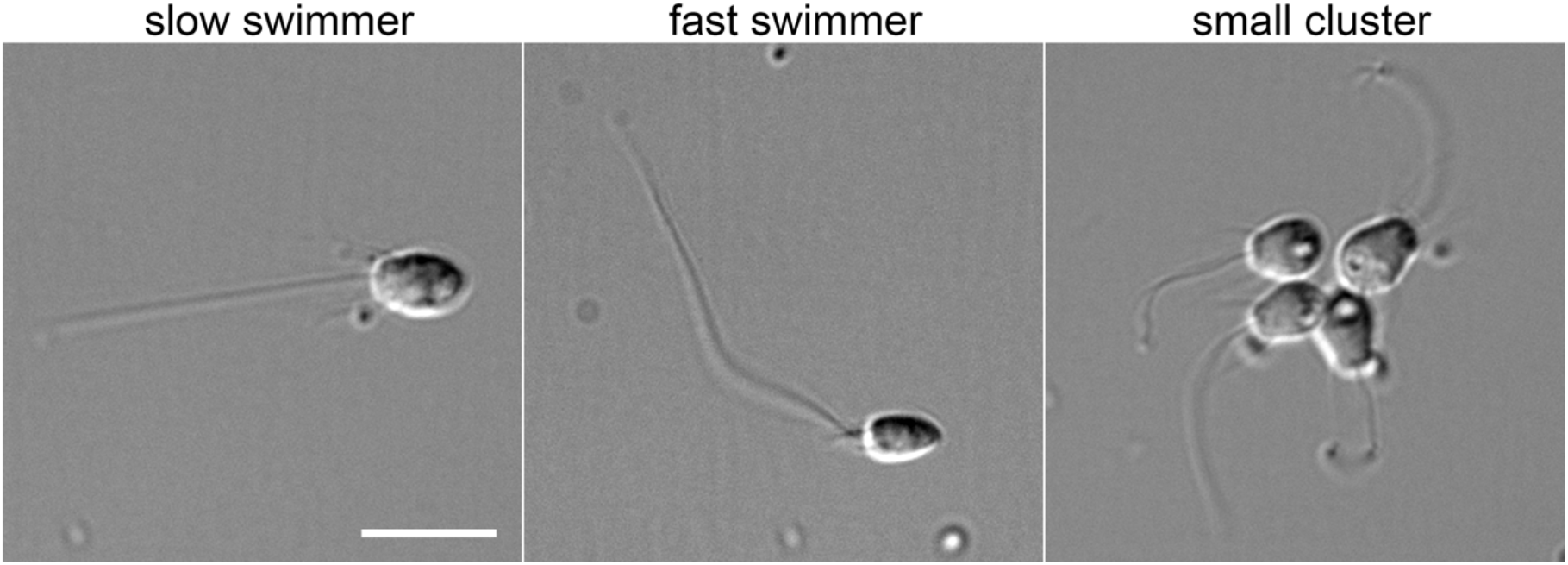
Morphology of a subset of *S. rosetta* cell types. Examples of choanoflagellate cell morphology from a single culture. Slow swimmers have a relatively large, rounded cell body, whereas fast swimmers have a smaller, pointier cell body, shorter collars, and a longer cilium. The cluster shown here is not a true “rosette”, but rather represents linked cells that likely recently divided. Cells were fixed with 2% glutaraldehyde and imaged at 60x magnification using differential interference contrast light microscopy. Scale bar, 10 µm.

**Figure 3-figure supplement 1.**
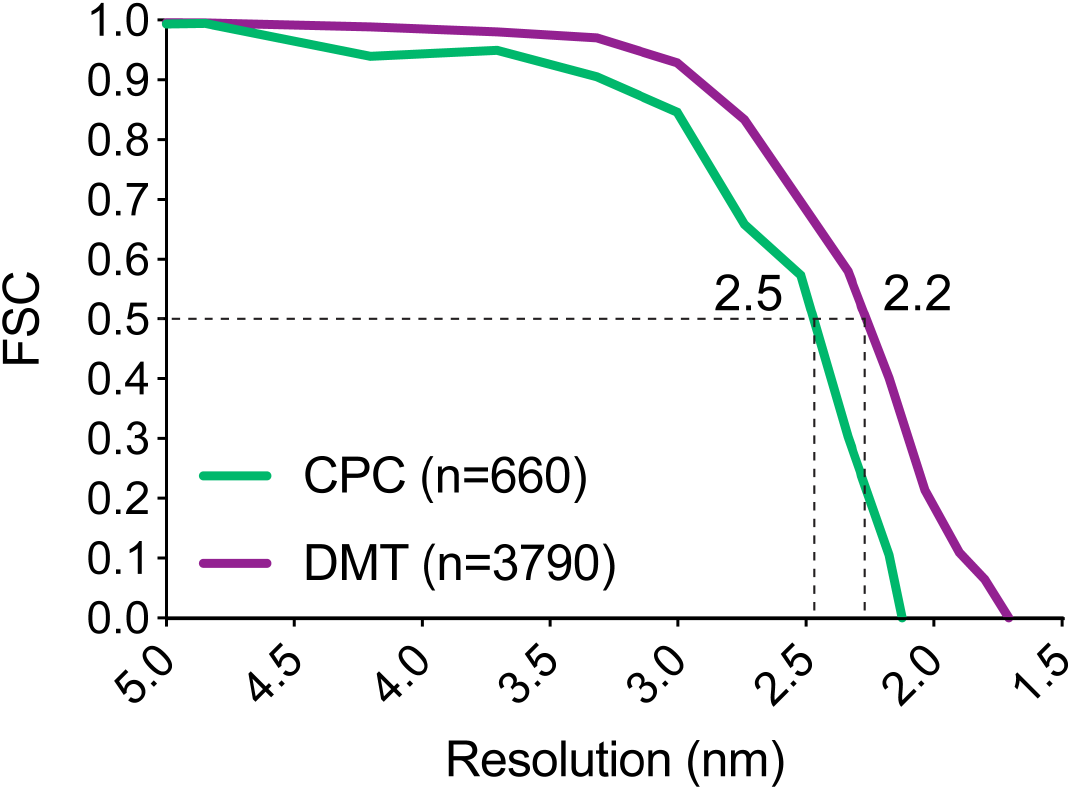
Resolution of averaged *S. rosetta* ciliary structures. Resolution estimates for the central pair complex average (CPC, green line) measured at the base of C1a and the 96-nm axonemal repeat (DMT, purple line) measured at the base of radial spoke 1 using the Fourier shell correlation method. Dotted lines and values indicate where the FSC curves intersect with the 0.5 criterion. Additional resolution information is listed in Table 1.

**Figure 3-figure supplement 2.**
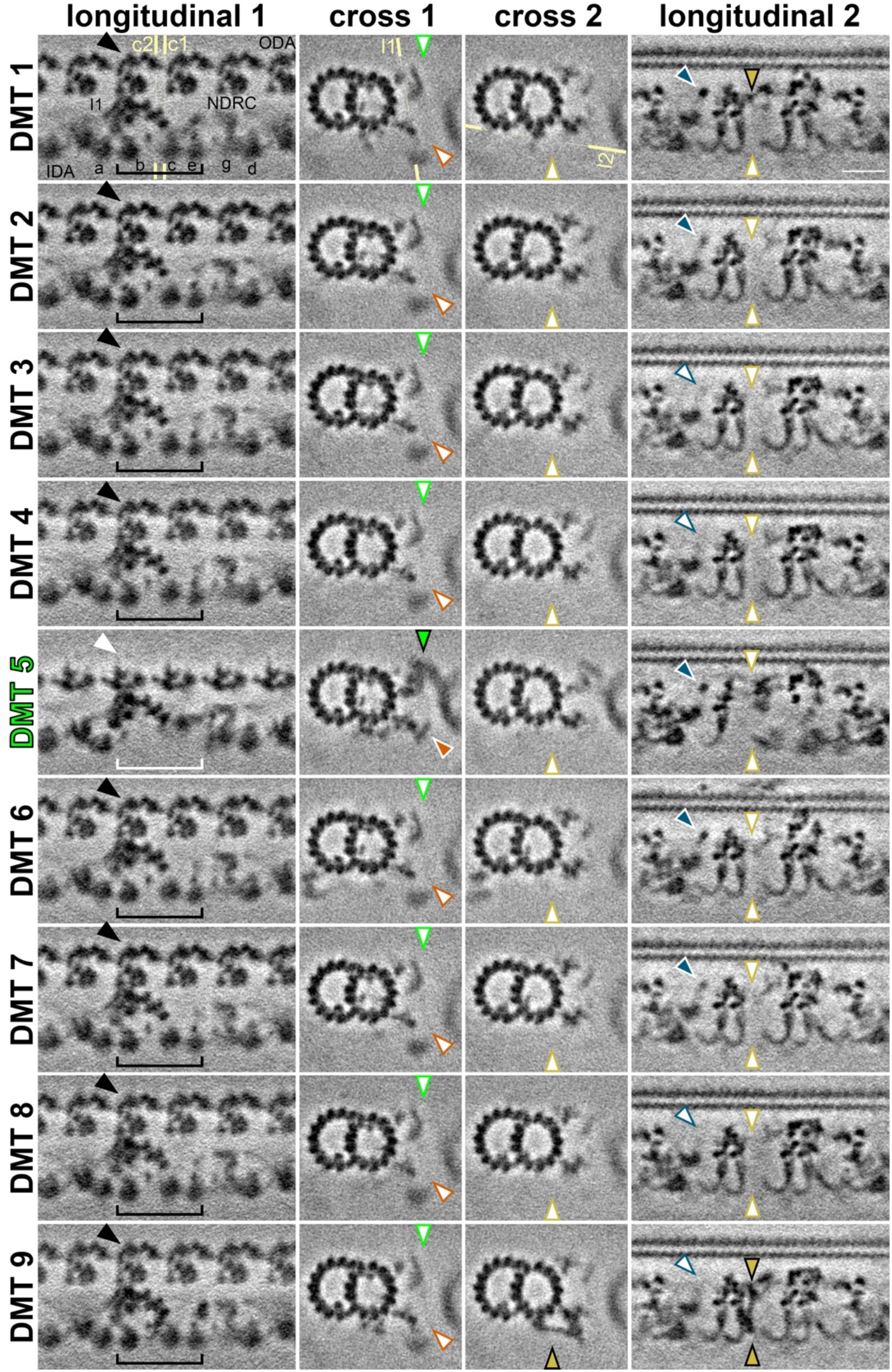
DMT-specific features in the *S. rosetta* cilium. Tomographic slices of the DMT-specific averages (from 54 tomograms, Table 1) show some features with asymmetric distribution. The column titled “longitudinal 1” highlights the absence of outer and inner dyneins from DMT 5 (white arrowhead/bracket indicates absence, black arrowhead/bracket indicates presence of ODAs and selected IDAs), which are replaced by a bridge between DMTs 5 and 6 (called o-SUB5-6, white arrowhead). The yellow lines indicate the positions of cross sections 1 and 2 (c1, c2), respectively. The column titled “cross 1” highlights the presence (colored arrowheads) of the o-SUB5-6 (green) and i-SUB5-6 (orange) on DMT 5, whereas these features are absent in all other DMTs (white arrowheads). The column titled “cross 2” highlights the presence (mustard arrowhead) of a unique connection between the A-tubule and the tail of IDAc only on DMT 9. The yellow line indicates the positions of slice shown in “longitudinal 2”. The column titled “longitudinal 2” shows a bottom-up view of the unique connection between the A-tubule and the tail of IDA c on DMT 9 with a similar density in the upper part of DMT 1 (mustard arrowheads), which are absent from all other DMTs. An additional density (blue arrowheads) is also present or partially present on DMTs 1, 2, 5, 6, and 7, but only weakly present or absent from DMTs 3, 4, 8, and 9 (white arrowheads), corresponding to the area just beneath the A-tubule hole described in Figure 4. Scale bar: 20 nm (upper right panel, applies to all).

**Figure 4-figure supplement 1.**
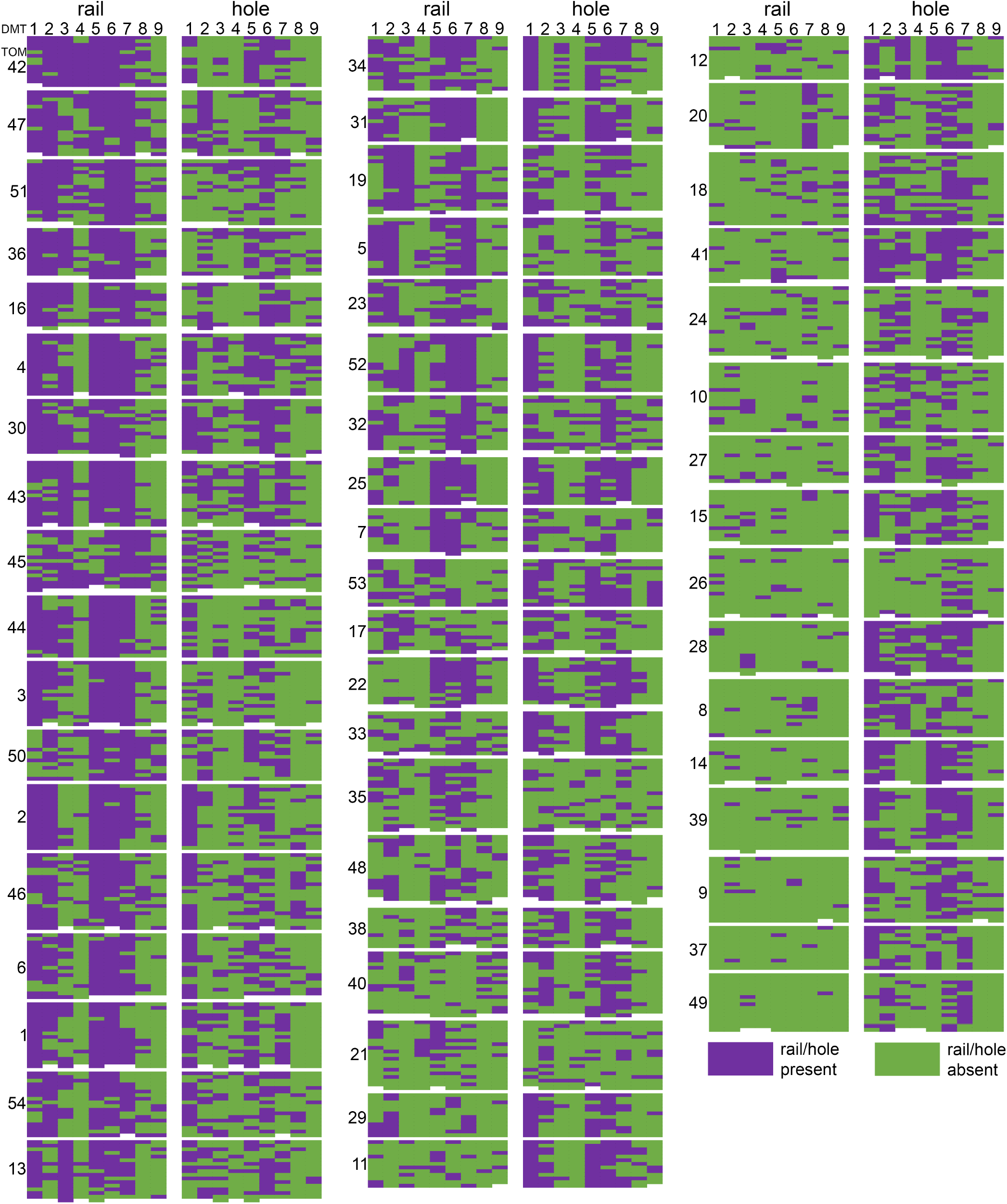
Distribution of rail-MIP and A-tubule hole classes by tomogram and doublet microtubule. Each tomogram (TOM **#**) is represented by a block in which the columns indicate microtubule doublet (DMT1-9), and the rows represent individual 96-nm subvolumes within those DMTs, listed from distal (top) to proximal (bottom). For each tomogram, the rail-MIP classification results are displayed on the left, and the A-tubule hole classes are displayed on the right. Purple indicates the presence of either the rail-MIP or the A-tubule hole, whereas green denotes their absence. Rail-MIPs seem to be strongly present in certain tomograms (left), and almost completely absent in others (right). Holes, on the other hand, exhibit a more scattered distribution. Both features show some DMT-specificity or -preference, but there is only partial overlap between the presence/absence of a rail-MIP and A-tubule hole in the same subvolume.

**Figure 4-figure supplement 2.**
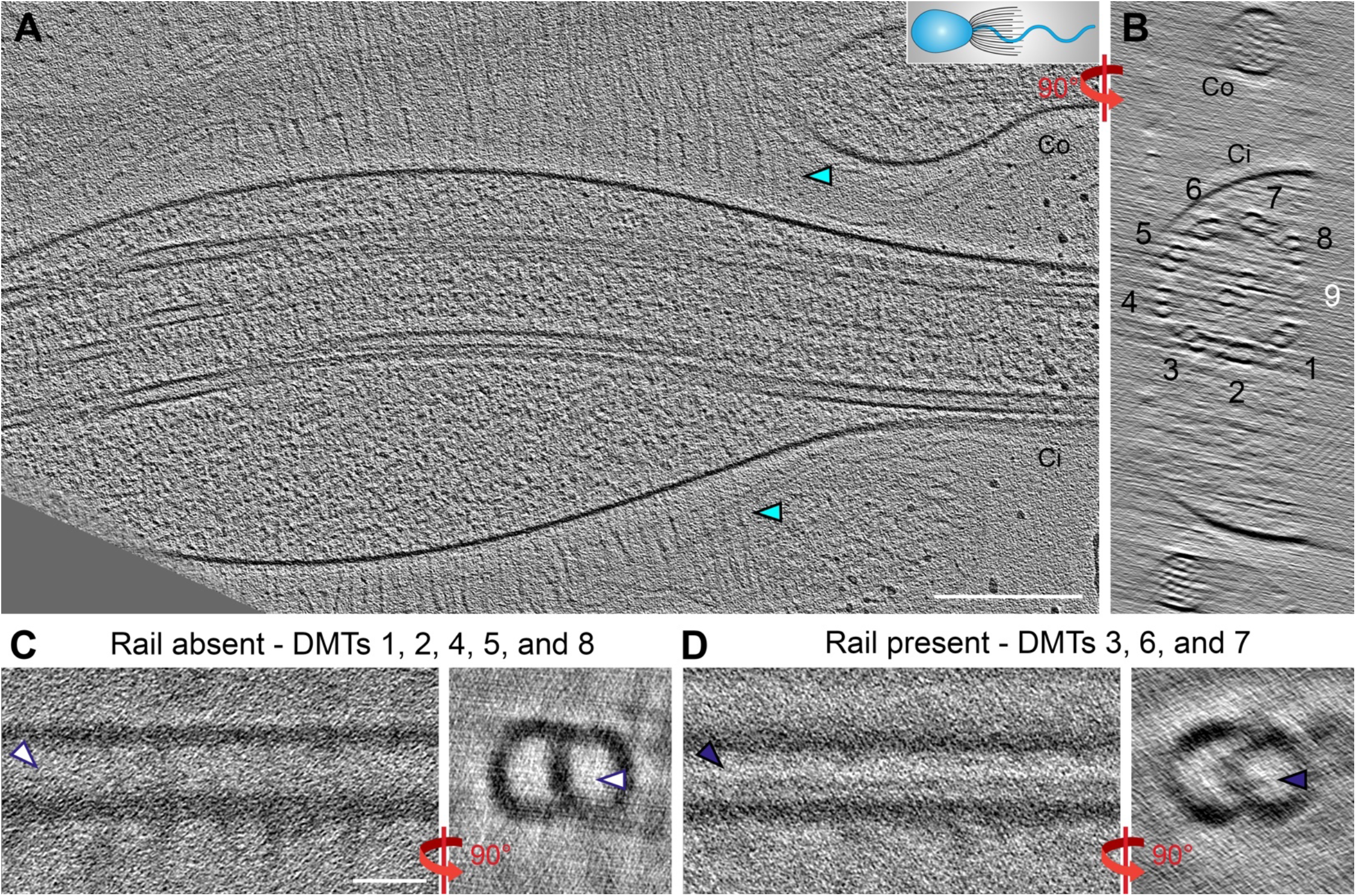
The rail-MIP is present in the proximal region of the *S. rosetta* cilium. **A**-**B**) Longitudinal (**A**) and cross-sectional (**B**) tomographic slices through a cryo-FIB milled *S. rosetta* cell, targeting the proximal region of the cilium close to the cell body. The cartoon in (**A**) denotes the cell’s orientation, with cell body to the left. Cyan arrowheads indicate vane filaments. Note: the ciliary membrane in this sample is slightly swollen, which is not unusual for cells that were actively swimming while they were plunge-frozen. DMT numbers are indicated, with DMT 9 in white to indicate its absence (the cryo-FIB-milling removed material from that side of the axoneme). **C**, **D**) Two subtomogram averages of the 96-nm axonemal repeat were calculated from the tomogram shown in (**A**, **B**): in (**C**), repeats from DMTs 1, 2, 4, 5, and 8 were combined, showing an average that lacks the rail-MIP (white arrowheads), and in (**D**), repeats from DMTs 3, 6, and 7 were combined, showing the rail-MIP (navy blue arrowheads). Note: the doublet-specific averages shown here are noisy due to limited number of averaged particles. Scale bars: 200 nm (A, applies also to B); 20 nm (C, applies also to D).

**Figure 4-figure supplement 3.**
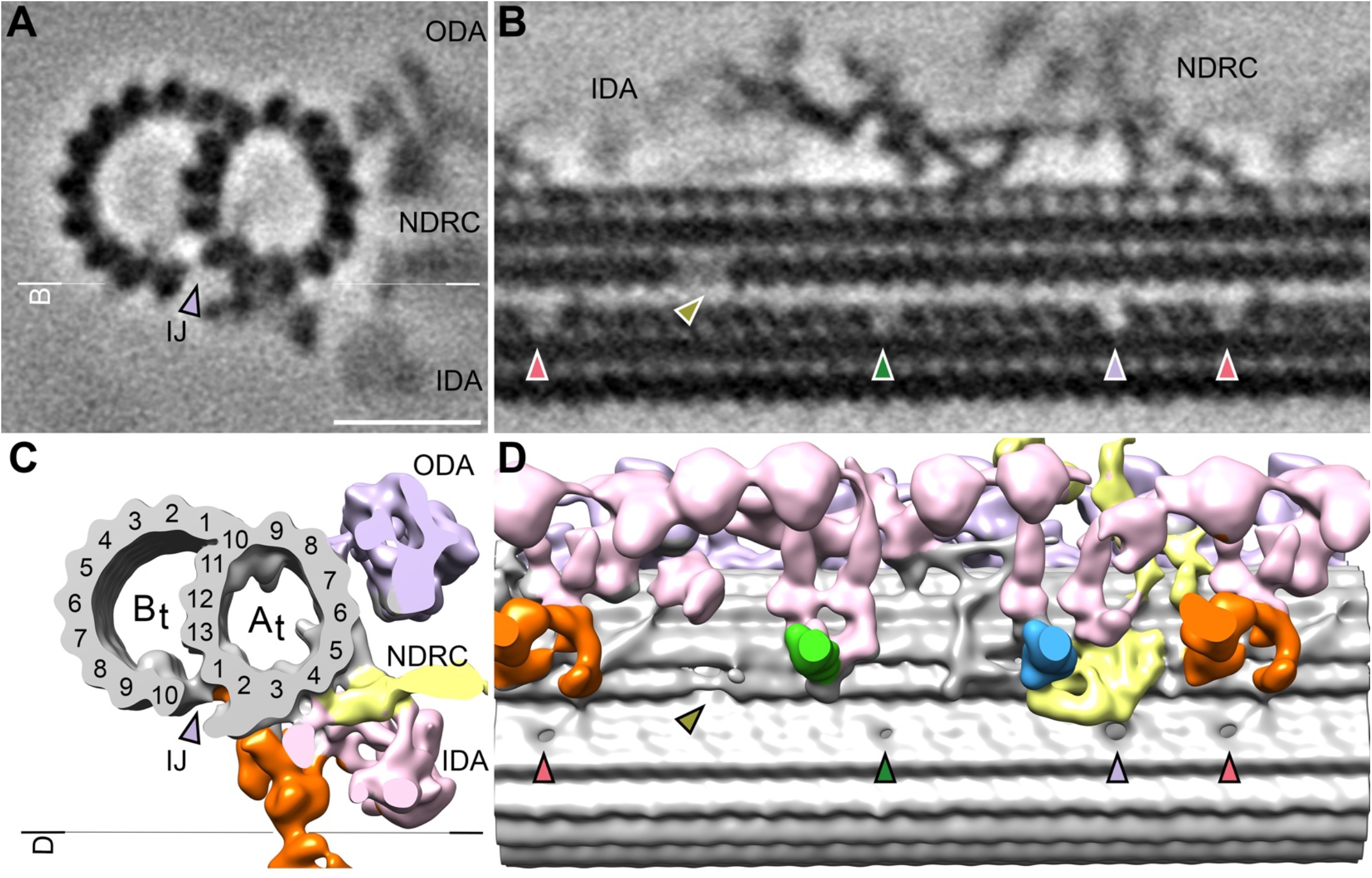
Additional holes in the doublet inner junction of the *S. rosetta* cilium. **A**-**B**) Tomographic slices through the subtomogram average of the *S. rosetta* 96-nm axonemal repeat shown in cross-section (**A**) and longitudinal section (**B**). The white line indicates the location of the section shown in (**B**). **C**-**D**) Isosurface renderings of the averages shown in (**A**, **B**). The black line indicates the view shown in (**D**). In all panels, the pale violet arrowheads indicate the previously reported hole near the N- DRC, the olive arrowheads indicate the A-tubule gap (see Figure 4 and Figure 4-figure supplement 1), and the green and pink arrows denote additional inner junction holes that have not been previously reported in other species. Other labels: IDA, inner dynein arms; NDRC, nexin-dynein regulatory complex; ODA, outer dynein arms. Scale bar: 20 nm (A, applies also to B).

**Figure 5-figure supplement 1.**
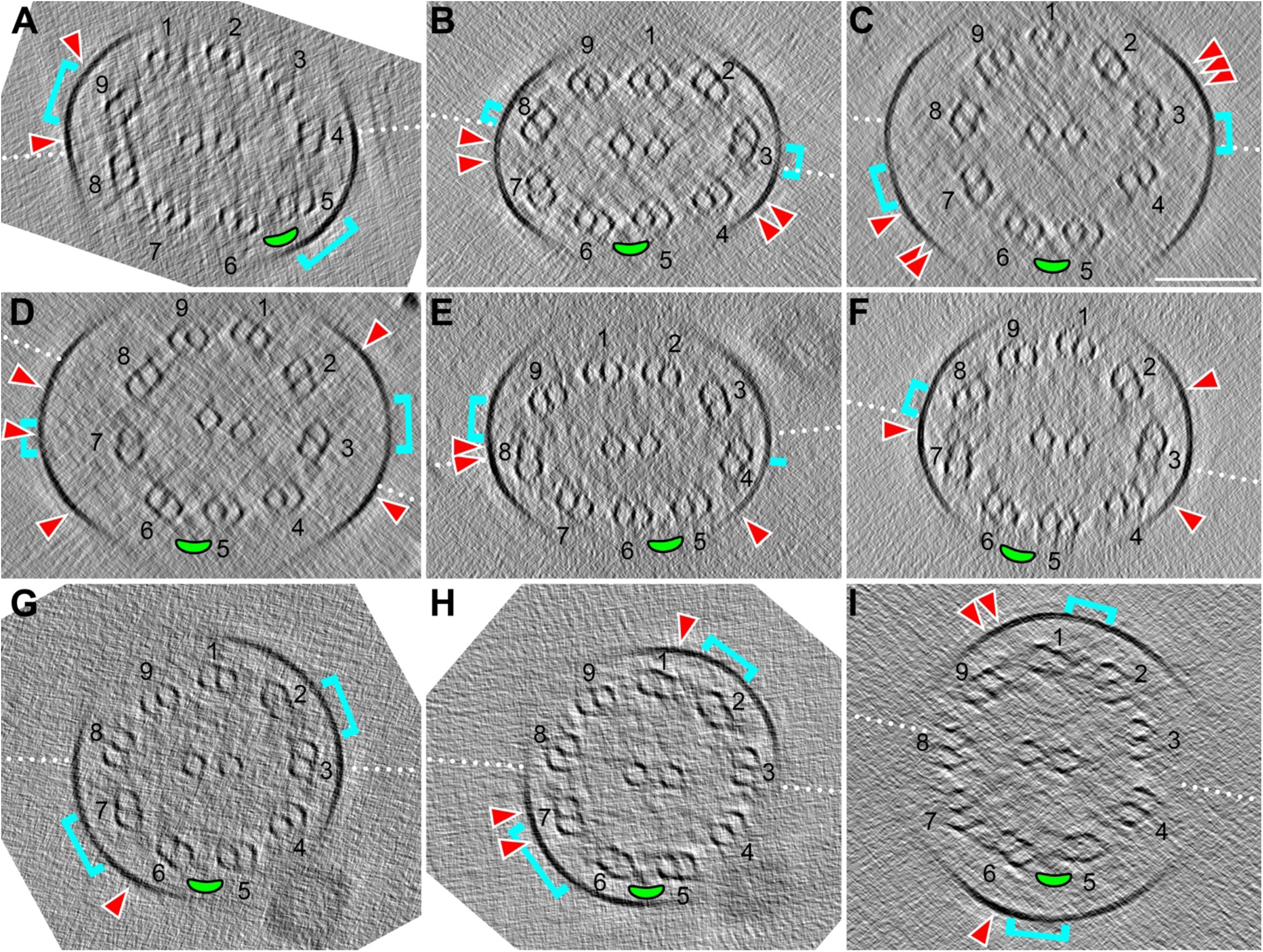
Relative orientation of CPC to DMTs and extraciliary features. **A**-**I**) Tomographic slices through individual *S. rosetta* cilia shown in cross-section. The white dotted lines indicate the CPC plane (i.e. the plane that contains both CPC microtubules). DMTs are labeled (1-9), with the 5-6 bridge highlighted with a green ellipse. The cyan brackets indicate the position on the membrane where vane filaments protrude, and the red arrowheads indicate the position of barbs (within ∼315 nm model depth). The plane of the CPC is consistently parallel to the 5-6 bridge, whereas the location of the vane and barbs varies slightly with respect to DMT and CPC position. Note: only two of the 56 analyzed tomograms showed an almost perpendicular orientation between the vane/barbs and the CPC/5-6 bridge plane (as in **I**). Scale bar: 100 nm (C, applies to all panels).

**Figure 5-figure supplement 2.**
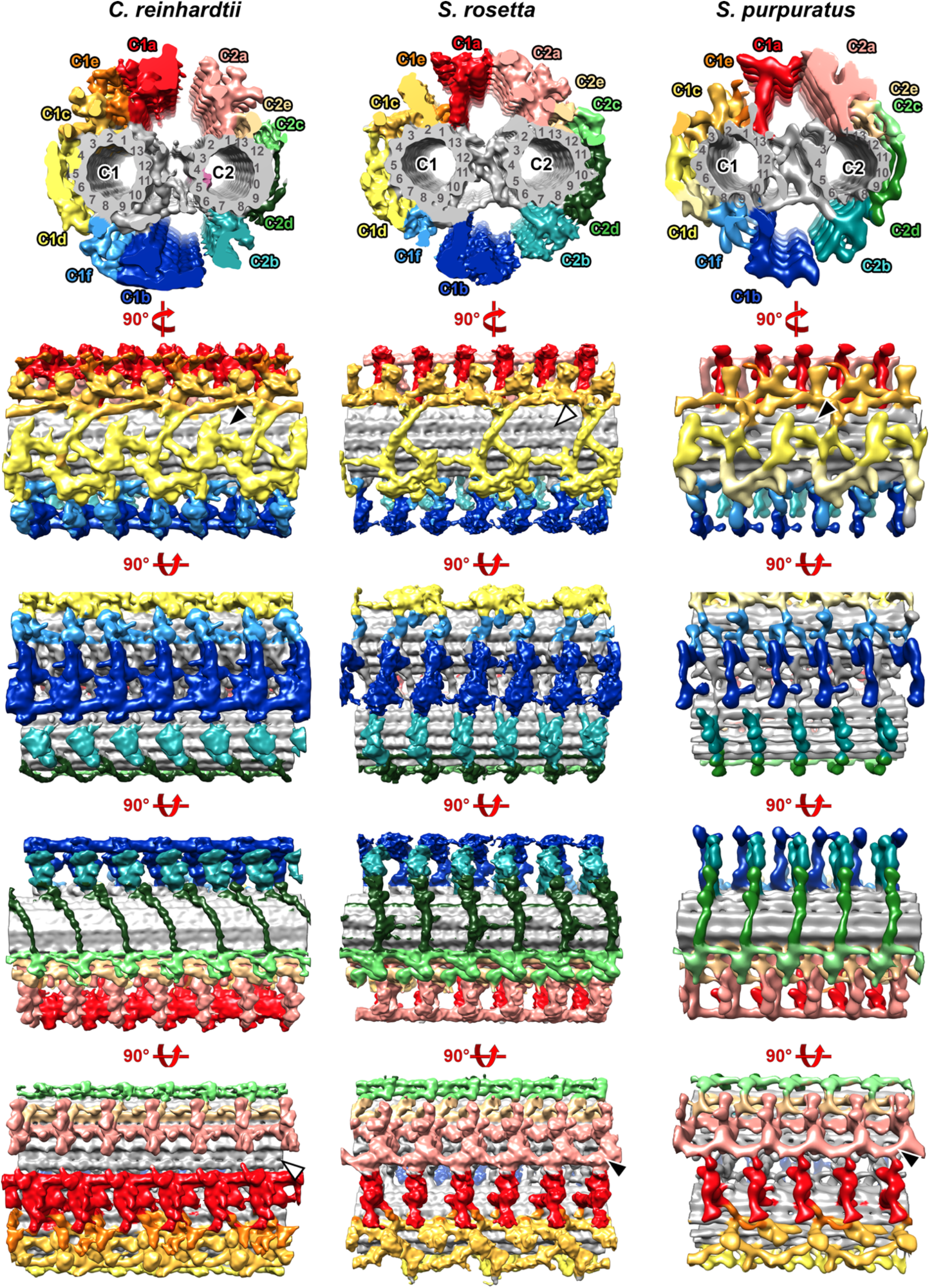
Evolutionary comparison of the central pair complex between *Chlamydomonas reinhardtii* (green alga)*, S. rosetta* (choanoflagellate), and *S. purpuratus* (sea urchin). Isosurface renderings are viewed in cross-section from proximal (top row) and rotated to generate the longitudinal views as indicated. Black and white arrowheads represent the presence and absence of indicated structures, respectively. The *C. reinhardtii* and *S. purpuratus* data were previously published (Carbajal-Gonzalez et al., 2013; Fu et al., 2019).

**Figure 6-figure supplement 1.**
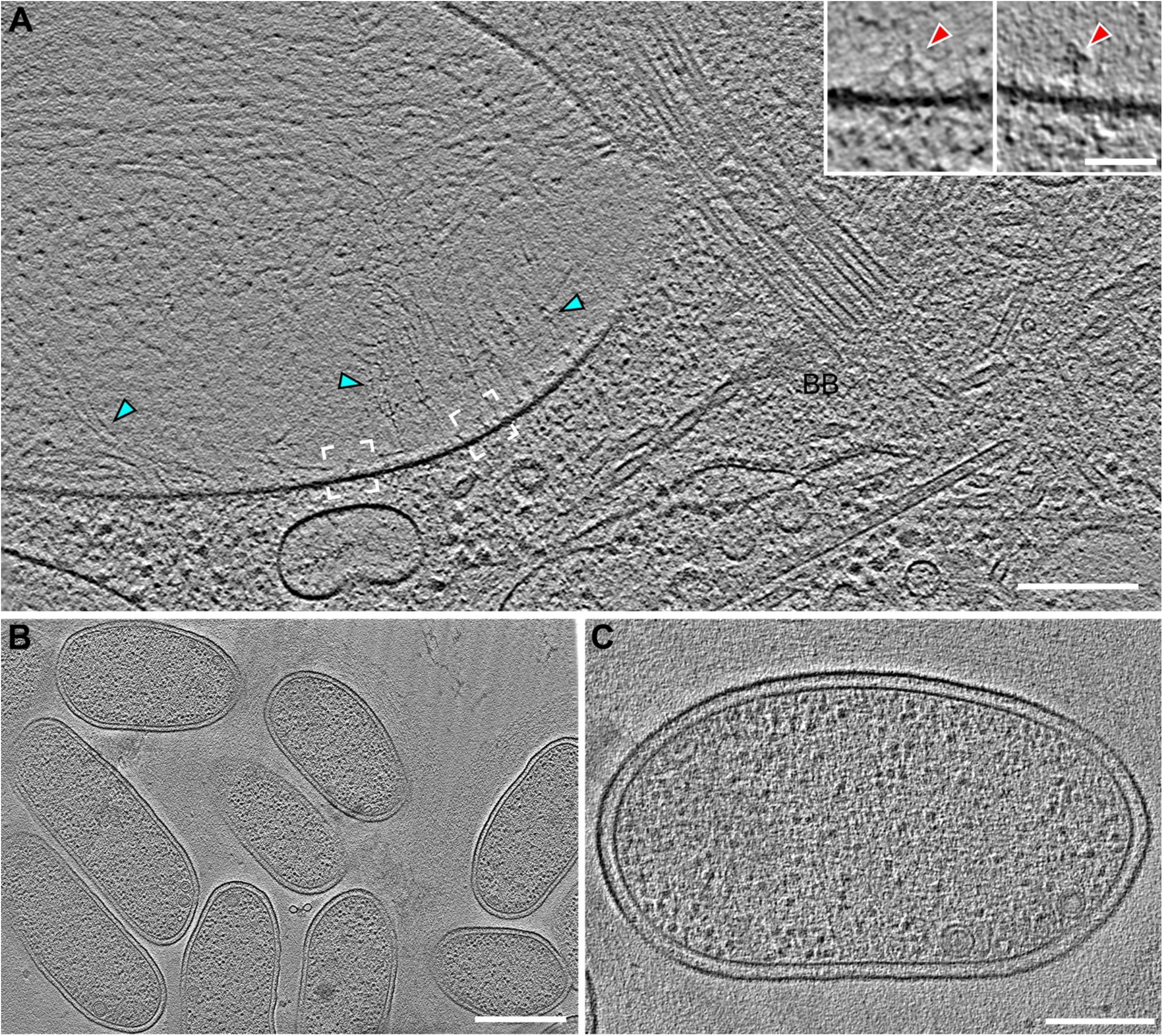
Vane filaments and barb-like structures are found in the ciliary pocket of *S. rosetta*, but not on or in the co-cultured bacterial prey cells. A) Tomographic slice through a cryo-FIB milled ciliary pocket/basal body region of a *S. rosetta* cell shows vane filaments (cyan arrowheads) attached to the plasma membrane in the ciliary pocket, which is separated from the ciliary membrane by a selective gating mechanism. White brackets denote positions of insets (left and right brackets correspond to left and right insets, respectively), which were slightly rotated to highlight the barb-like structures within the plasma membrane. B, C) Representative tomographic slices through *E pacifica* bacteria, which are co-cultured with *S. rosetta* as a food source, showing smooth membranes that lack vane filaments and barb structures. Scale bars: 200 nm (A); 50 nm (A inset); 500 nm (B); 200 nm (C).

## References

Afzelius, B. (1959). Electron microscopy of the sperm tail; results obtained with a new fixative. J Biophys Biochem Cytol 5, 269–278.

Barber, C.F., Heuser, T., Carbajal-Gonzalez, B.I., Botchkarev, V.V., Jr., and Nicastro, D. (2012). Three-dimensional structure of the radial spokes reveals heterogeneity and interactions with dyneins in Chlamydomonas flagella. Mol Biol Cell 23, 111–120.

Blake, J.R., and Sleigh, M.A. (1974). Mechanics of ciliary locomotion. Biol Rev Camb Philos Soc 49, 85–125.

Bloodgood, R.A. (2010). Sensory reception is an attribute of both primary cilia and motile cilia. J Cell Sci 123, 505–509.

Booth, D.S., and King, N. (2020). Genome editing enables reverse genetics of multicellular development in the choanoflagellate Salpingoeca rosetta. Elife 9.

Booth, D.S., Szmidt-Middleton, H., and King, N. (2018). Transfection of choanoflagellates illuminates their cell biology and the ancestry of animal septins. Mol Biol Cell 29, 3026–3038.

Bouck, G.B. (1971). The structure, origin, isolation, and composition of the tubular mastigonemes of the Ochromas flagellum. J Cell Biol 50, 362–384.

Brunet, T., and King, N. (2017). The Origin of Animal Multicellularity and Cell Differentiation. Dev Cell 43, 124–140.

Burki, F. (2014). The eukaryotic tree of life from a global phylogenomic perspective. Cold Spring Harb Perspect Biol 6, a016147.

Carbajal-Gonzalez, B.I., Heuser, T., Fu, X., Lin, J., Smith, B.W., Mitchell, D.R., and Nicastro, D. (2013). Conserved structural motifs in the central pair complex of eukaryotic flagella. Cytoskeleton (Hoboken) 70, 101–120.

Carr, M., Leadbeater, B.S., Hassan, R., Nelson, M., and Baldauf, S.L. (2008). Molecular phylogeny of choanoflagellates, the sister group to Metazoa. Proc Natl Acad Sci U S A 105, 16641–16646.

Cavalier-Smith, T. (2002). The phagotrophic origin of eukaryotes and phylogenetic classification of Protozoa. Int J Syst Evol Microbiol 52, 297–354.

Danev, R., Buijsse, B., Khoshouei, M., Plitzko, J.M., and Baumeister, W. (2014). Volta potential phase plate for in-focus phase contrast transmission electron microscopy. Proc Natl Acad Sci U S A 111, 15635–15640.

Dayel, M.J., Alegado, R.A., Fairclough, S.R., Levin, T.C., Nichols, S.A., McDonald, K., and King, N. (2011). Cell differentiation and morphogenesis in the colony-forming choanoflagellate Salpingoeca rosetta. Dev Biol 357, 73–82.

Dayel, M.J., and King, N. (2014). Prey capture and phagocytosis in the choanoflagellate Salpingoeca rosetta. PLoS One 9, e95577.

de Souza, W., and Souto-Padron, T. (1980). The paraxial structure of the flagellum of trypanosomatidae. J Parasitol 66, 229–236.

Dobro, M.J., Oikonomou, C.M., Piper, A., Cohen, J., Guo, K., Jensen, T., Tadayon, J., Donermeyer, J., Park, Y., Solis, B.A., et al. (2017). Uncharacterized Bacterial Structures Revealed by Electron Cryotomography. J Bacteriol 199.

Dymek, E.E., Lin, J., Fu, G., Porter, M.E., Nicastro, D., and Smith, E.F. (2019). PACRG and FAP20 form the inner junction of axonemal doublet microtubules and regulate ciliary motility. Mol Biol Cell 30, 1805–1816.

Fawcett, D.W. (1954). The study of epithelial cilia and sperm flagella with the electron microscope. Laryngoscope 64, 557–567.

Fu, G., Wang, Q., Phan, N., Urbanska, P., Joachimiak, E., Lin, J., Wloga, D., and Nicastro, D. (2018). The I1 dynein-associated tether and tether head complex is a conserved regulator of ciliary motility. Mol Biol Cell 29, 1048–1059.

Fu, G., Zhao, L., Dymek, E., Hou, Y., Song, K., Phan, N., Shang, Z., Smith, E.F., Witman, G.B., and Nicastro, D. (2019). Structural organization of the C1a-e-c supercomplex within the ciliary central apparatus. J Cell Biol 218, 4236–4251.

Hagen, W.J.H., Wan, W., and Briggs, J.A.G. (2017). Implementation of a cryo-electron tomography tilt-scheme optimized for high resolution subtomogram averaging. J Struct Biol 197, 191–198.

Han, L., Rao, Q., Yang, R., Wang, Y., Chai, P., Xiong, Y., and Zhang, K. (2022). Cryo-EM structure of an active central apparatus. bioRxiv, 2022.2001.2023.477438.

Heumann, J.M., Hoenger, A., and Mastronarde, D.N. (2011). Clustering and variance maps for cryo-electron tomography using wedge-masked differences. J Struct Biol 175, 288–299.

Hibberd, D.J. (1975). Observations on the ultrastructure of the choanoflagellate Codosiga botrytis (Ehr.) Saville-Kent with special reference to the flagellar apparatus. J Cell Sci 17, 191–219.

Hoops, H.J., and Witman, G.B. (1983). Outer doublet heterogeneity reveals structural polarity related to beat direction in Chlamydomonas flagella. J Cell Biol 97, 902–908.

Hyams, J.S. (1982). The Euglena paraflagellar rod: structure, relationship to other flagellar components and preliminary biochemical characterization. J Cell Sci 55, 199–210.

Iancu, C.V., Tivol, W.F., Schooler, J.B., Dias, D.P., Henderson, G.P., Murphy, G.E., Wright, E.R., Li, Z., Yu, Z., Briegel, A., et al. (2006). Electron cryotomography sample preparation using the Vitrobot. Nat Protoc 1, 2813–2819.

Ichikawa, M., Liu, D., Kastritis, P.L., Basu, K., Hsu, T.C., Yang, S., and Bui, K.H. (2017). Subnanometre-resolution structure of the doublet microtubule reveals new classes of microtubule-associated proteins. Nat Commun 8, 15035.

Imhof, S., Zhang, J., Wang, H., Bui, K.H., Nguyen, H., Atanasov, I., Hui, W.H., Yang, S.K., Zhou, Z.H., and Hill, K.L. (2019). Cryo electron tomography with volta phase plate reveals novel structural foundations of the 96-nm axonemal repeat in the pathogen Trypanosoma brucei. Elife 8.

Irons, M.J., and Clermont, Y. (1982a). Formation of the outer dense fibers during spermiogenesis in the rat. Anat Rec 202, 463–471.

Irons, M.J., and Clermont, Y. (1982b). Kinetics of fibrous sheath formation in the rat spermatid. Am J Anat 165, 121–130.

Karpov, S.A. (2016). Flagellar apparatus structure of choanoflagellates. Cilia 5, 11.

Karpov, S.A., and Leadbeater, B.S.C. (1998). Cytoskeleton Structure and Composition in Choanoflagellates. J Euk Microbiol 45, 361–367.

Khalifa, A.A.Z., Ichikawa, M., Dai, D., Kubo, S., Black, C.S., Peri, K., McAlear, T.S., Veyron, S., Yang, S.K., Vargas, J., et al. (2020). The inner junction complex of the cilia is an interaction hub that involves tubulin post-translational modifications. Elife 9.

King, N. (2004). The unicellular ancestry of animal development. Dev Cell 7, 313–325.

Kirima, J., and Oiwa, K. (2018). Flagellar-associated Protein FAP85 Is a Microtubule Inner Protein That Stabilizes Microtubules. Cell Struct Funct 43, 1–14.

Kollmar, M. (2016). Fine-Tuning Motile Cilia and Flagella: Evolution of the Dynein Motor Proteins from Plants to Humans at High Resolution. Molecular biology and evolution 33, 3249–3267.

Kremer, J.R., Mastronarde, D.N., and McIntosh, J.R. (1996). Computer visualization of three-dimensional image data using IMOD. J Struct Biol 116, 71–76.

Leadbeater, B. (2006). The ’mystery’ of the flagellar vane in choanoflagellates. Nova Hedwigia, 213-223. Leadbeater, B.S.C. (2015). The Choanoflagellates (Cambridge University Press).

Leung, M.R., Roelofs, M.C., Ravi, R.T., Maitan, P., Henning, H., Zhang, M., Bromfield, E.G., Howes, S.C., Gadella, B.M., Bloomfield-Gadêlha, H., et al. (2021). The multi-scale architecture of mammalian sperm flagella and implications for ciliary motility. The EMBO Journal 40, e107410.

Li, S., Fernandez, J.J., Fabritius, A.S., Agard, D.A., and Winey, M. (2022). Electron cryo-tomography structure of axonemal doublet microtubule from Tetrahymena thermophila. Life Sci Alliance 5.

Lin, J., Heuser, T., Carbajal-Gonzalez, B.I., Song, K., and Nicastro, D. (2012a). The structural heterogeneity of radial spokes in cilia and flagella is conserved. Cytoskeleton (Hoboken) 69, 88–100.

Lin, J., Heuser, T., Song, K., Fu, X., and Nicastro, D. (2012b). One of the nine doublet microtubules of eukaryotic flagella exhibits unique and partially conserved structures. PLoS One 7, e46494.

Lin, J., and Nicastro, D. (2018). Asymmetric distribution and spatial switching of dynein activity generates ciliary motility. Science 360.

Lin, J., Yin, W., Smith, M.C., Song, K., Leigh, M.W., Zariwala, M.A., Knowles, M.R., Ostrowski, L.E., and Nicastro, D. (2014). Cryo-electron tomography reveals ciliary defects underlying human RSPH1 primary ciliary dyskinesia. Nat Commun 5, 5727.

Liu, P., Lou, X., Wingfield, J.L., Lin, J., Nicastro, D., and Lechtreck, K. (2020). Chlamydomonas PKD2 organizes mastigonemes, hair-like glycoprotein polymers on cilia. J Cell Biol 219.

Ma, M., Stoyanova, M., Rademacher, G., Dutcher, S.K., Brown, A., and Zhang, R. (2019). Structure of the Decorated Ciliary Doublet Microtubule. Cell 179, 909–922 e912.

Mah, J.L., Christensen-Dalsgaard, K.K., and Leys, S.P. (2014). Choanoflagellate and choanocyte collar- flagellar systems and the assumption of homology. Evol Dev 16, 25–37.

Maheshwari, A., Obbineni, J.M., Bui, K.H., Shibata, K., Toyoshima, Y.Y., and Ishikawa, T. (2015). alpha- and beta-Tubulin Lattice of the Axonemal Microtubule Doublet and Binding Proteins Revealed by Single Particle Cryo-Electron Microscopy and Tomography. Structure 23, 1584–1595.

Marko, M., Hsieh, C., Schalek, R., Frank, J., and Mannella, C. (2007). Focused-ion-beam thinning of frozen-hydrated biological specimens for cryo-electron microscopy. Nat Methods 4, 215–217.

Mastronarde, D.N. (2005). Automated electron microscope tomography using robust prediction of specimen movements. J Struct Biol 152, 36–51.

Matriano, D.M., Alegado, R.A., and Conaco, C. (2021). Detection of horizontal gene transfer in the genome of the choanoflagellate Salpingoeca rosetta. Sci Rep 11, 5993.

McIntosh, R., Nicastro, D., and Mastronarde, D. (2005). New views of cells in 3D: an introduction to electron tomography. Trends Cell Biol 15, 43–51.

Mehl, D., and Reiswig, H.M. (1991). The presence of flagellar vanes in choanomeres of Porifera and their possible phylogenetic implications. Journal of Zoological Systematics and Evolutionary Research 29, 312–319.

Mencarelli, C., Lupetti, P., and Dallai, R. (2008). New insights into the cell biology of insect axonemes. Int Rev Cell Mol Biol 268, 95–145.

Mitchell, D.R. (2004). Speculations on the evolution of 9+2 organelles and the role of central pair microtubules. Biology of the cell 96, 691–696.

Mitchell, D.R. (2007). The evolution of eukaryotic cilia and flagella as motile and sensory organelles. Adv Exp Med Biol 607, 130–140.

Nakamura, S., Tanaka, G., Maeda, T., Kamiya, R., Matsunaga, T., and Nikaido, O. (1996). Assembly and function of Chlamydomonas flagellar mastigonemes as probed with a monoclonal antibody. J Cell Sci 109 ( *Pt 1*), 57–62.

Nicastro, D., Fu, X., Heuser, T., Tso, A., Porter, M.E., and Linck, R.W. (2011). Cryo-electron tomography reveals conserved features of doublet microtubules in flagella. Proc Natl Acad Sci U S A 108, E845–853.

Nicastro, D., Schwartz, C., Pierson, J., Gaudette, R., Porter, M.E., and McIntosh, J.R. (2006). The molecular architecture of axonemes revealed by cryoelectron tomography. Science 313, 944–948.

Nielsen, L.T., Asadzadeh, S.S., Dolger, J., Walther, J.H., Kiorboe, T., and Andersen, A. (2017). Hydrodynamics of microbial filter feeding. Proc Natl Acad Sci U S A 114, 9373–9378.

Omoto, C.K., Gibbons, I.R., Kamiya, R., Shingyoji, C., Takahashi, K., and Witman, G.B. (1999). Rotation of the central pair microtubules in eukaryotic flagella. Mol Biol Cell 10, 1–4.

Owa, M., Uchihashi, T., Yanagisawa, H.A., Yamano, T., Iguchi, H., Fukuzawa, H., Wakabayashi, K.I., Ando, T., and Kikkawa, M. (2019). Inner lumen proteins stabilize doublet microtubules in cilia and flagella. Nat Commun 10, 1143.

Pazour, G.J., Agrin, N., Leszyk, J., and Witman, G.B. (2005). Proteomic analysis of a eukaryotic cilium. J Cell Biol 170, 103–113.

Pettersen, E.F., Goddard, T.D., Huang, C.C., Couch, G.S., Greenblatt, D.M., Meng, E.C., and Ferrin, T.E. (2004). UCSF Chimera--a visualization system for exploratory research and analysis. J Comput Chem 25, 1605–1612.

Pettitt, M., Orme, B., Blake, J.R., and Leadbeater, B.S. (2002). The hydrodynamics of filter feeding in choanoflagellates. European Journal of Protistology 38, 313–332.

Pigino, G., Bui, K.H., Maheshwari, A., Lupetti, P., Diener, D., and Ishikawa, T. (2011). Cryoelectron tomography of radial spokes in cilia and flagella. J Cell Biol 195, 673–687.

Pigino, G., Maheshwari, A., Bui, K.H., Shingyoji, C., Kamimura, S., and Ishikawa, T. (2012). Comparative structural analysis of eukaryotic flagella and cilia from Chlamydomonas, Tetrahymena, and sea urchins. J Struct Biol 178, 199–206.

Poghosyan, E., Iacovache, I., Faltova, L., Leitner, A., Yang, P., Diener, D.R., Aebersold, R., Zuber, B., and Ishikawa, T. (2020). The structure and symmetry of the radial spoke protein complex in Chlamydomonas flagella. J Cell Sci 133.

Porter, M.E., and Sale, W.S. (2000). The 9 + 2 axoneme anchors multiple inner arm dyneins and a network of kinases and phosphatases that control motility. J Cell Biol 151, F37–42.

Portman, N., and Gull, K. (2010). The paraflagellar rod of kinetoplastid parasites: from structure to components and function. Int J Parasitol 40, 135–148.

Pozdnyakov, I., R, Sokolova, A., M, Ereskovsky, A., and Karpov, S., A (2017). Kinetid structure of choanoflagellates and choanocytes of sponges does not support their close relationship. Protistology 11, 248–264.

Reiter, J.F., and Leroux, M.R. (2017). Genes and molecular pathways underpinning ciliopathies. Nat Rev Mol Cell Biol 18, 533–547.

Ruiz-Trillo, I., Roger, A.J., Burger, G., Gray, M.W., and Lang, B.F. (2008). A phylogenomic investigation into the origin of metazoa. Molecular biology and evolution 25, 664–672.

Schaffer, M., Mahamid, J., Engel, B.D., Laugks, T., Baumeister, W., and Plitzko, J.M. (2017). Optimized cryo-focused ion beam sample preparation aimed at in situ structural studies of membrane proteins. J Struct Biol 197, 73–82.

Schwartz, C.L., Heumann, J.M., Dawson, S.C., and Hoenger, A. (2012). A detailed, hierarchical study of Giardia lamblia’s ventral disc reveals novel microtubule-associated protein complexes. PLoS One 7, e43783.

Sigg, M.A., Menchen, T., Lee, C., Johnson, J., Jungnickel, M.K., Choksi, S.P., Garcia, G., 3rd, Busengdal, H., Dougherty, G.W., Pennekamp, P., et al. (2017). Evolutionary Proteomics Uncovers Ancient Associations of Cilia with Signaling Pathways. Dev Cell 43, 744–762 e711.

Smith, E.F., and Yang, P. (2004). The radial spokes and central apparatus: mechano-chemical transducers that regulate flagellar motility. Cell Motil Cytoskeleton 57, 8–17.

Song, K., Shang, Z., Fu, X., Lou, X., Grigorieff, N., and Nicastro, D. (2020). In situ structure determination at nanometer resolution using TYGRESS. Nat Methods 17, 201–208.

Steenkamp, E.T., Wright, J., and Baldauf, S.L. (2006). The protistan origins of animals and fungi. Molecular biology and evolution 23, 93–106.

Takeda, S., and Narita, K. (2012). Structure and function of vertebrate cilia, towards a new taxonomy. Differentiation 83, S4–11.

Wang, X., Fu, Y., Beatty, W.L., Ma, M., Brown, A., Sibley, L.D., and Zhang, R. (2021). Cryo-EM structure of cortical microtubules from human parasite Toxoplasma gondii identifies their microtubule inner proteins. Nat Commun 12, 3065.

Wetzel, L.A., Levin, T.C., Hulett, R.E., Chan, D., King, G.A., Aldayafleh, R., Booth, D.S., Sigg, M.A., and King, N. (2018). Predicted glycosyltransferases promote development and prevent spurious cell clumping in the choanoflagellate S. rosetta. Elife 7.

Yamaguchi, H., Oda, T., Kikkawa, M., and Takeda, H. (2018). Systematic studies of all PIH proteins in zebrafish reveal their distinct roles in axonemal dynein assembly. Elife 7.

Yubuki, N., Huang, S.S., and Leander, B.S. (2016). Comparative Ultrastructure of Fornicate Excavates, Including a Novel Free-living Relative of Diplomonads: Aduncisulcus paluster gen. et sp. nov. Protist 167, 584–596.

Zhao, Y., Pinskey, J., Lin, J., Yin, W., Sears, P.R., Daniels, L.A., Zariwala, M.A., Knowles, M.R., Ostrowski, L.E., and Nicastro, D. (2021). Structural insights into the cause of human RSPH4A primary ciliary dyskinesia. Mol Biol Cell 32, 1202–1209.

Zheng, W., Li, F., Ding, Z., Liu, H., Zhu, L., Xu, C., Li, J., Gao, Q., Wang, Y., Fu, Z., et al. (2021). Distinct architecture and composition of mouse axonemal radial spoke head revealed by cryo-EM. Proceedings of the National Academy of Sciences 118, e2021180118.

Zhu, X., Liu, Y., and Yang, P. (2017). Radial Spokes-A Snapshot of the Motility Regulation, Assembly, and Evolution of Cilia and Flagella. Cold Spring Harb Perspect Biol 9.

